# Inhibition of protein N-myristoylation blocks *Plasmodium falciparum* intraerythrocytic development, egress, and invasion

**DOI:** 10.1101/2020.12.16.423054

**Authors:** Anja C. Schlott, Ellen Knuepfer, Judith L. Green, Philip Hobson, Aaron J. Borg, Julia Morales-Sanfrutos, Abigail J. Perrin, Ambrosius P. Snijders, Edward W Tate, Anthony A. Holder

## Abstract

We have combined chemical biology and genetic modification approaches to investigate the importance of protein myristoylation in the human malaria parasite, *Plasmodium falciparum*. Parasite treatment during schizogony in the last ten to fifteen hours of the erythrocytic cycle with IMP-1002, an inhibitor of *N*-myristoyl transferase (NMT), led to a significant blockade in parasite egress from the infected erythrocyte. Two rhoptry proteins were mislocalized in the cell, suggesting that rhoptry function is disrupted. We identified sixteen NMT substrates for which myristoylation was significantly reduced by NMT inhibitor treatment, and of these, six proteins were substantially reduced in abundance. In a viability screen, we showed that for four of these proteins replacement of the N-terminal glycine with alanine to prevent myristoylation had a substantial effect on parasite fitness. In detailed studies of one NMT substrate, glideosome associated protein 45 (GAP45), loss of myristoylation had no impact on protein location or glideosome assembly, in contrast to the disruption caused by GAP45 gene deletion, but GAP45 myristoylation was essential for erythrocyte invasion. Therefore, there are at least three mechanisms by which inhibition of NMT can disrupt parasite development and growth: early in parasite development, leading to the inhibition of schizogony and formation of ‘pseudoschizonts’, which has been described previously; at the end of schizogony, with disruption of rhoptry formation, merozoite development and egress from the infected erythrocyte; and at invasion, when impairment of motor complex function prevents invasion of new erythrocytes. These results underline the importance of *P. falciparum* NMT as a drug target because of the pleiotropic effect of its inhibition.

## INTRODUCTION

The malarial parasite asexual blood stage is largely intra-erythrocytic as the parasite invades, develops and proliferates within red blood cells (RBCs) over a period of approximately 45 – 48 hours in the case of *Plasmodium falciparum*, the most lethal human parasite. Following invasion by the extracellular merozoite the parasite profoundly modifies the RBC, growing through ring and trophozoite stages and then starting multiple rounds of nuclear division around 30 hours after invasion, resulting in schizont formation. Coincident with nuclear division, the parasite constructs a series of subcellular membranous structures that will form the inner membrane complex (IMC) and apical organelles such as the rhoptries and micronemes of the nascent 20 - 30 daughter merozoites. At the end of schizogony the multinucleate coenocyte undergoes cytokinesis that draws the parasite plasma membrane (PM) around each of the developing progeny to form highly polarised merozoites, each with its own nucleus, a surface pellicle comprised of PM and IMC, and apical organelles for subsequent invasion and modification of a new RBC. Completion of this process is followed by lysis of the infected RBC and egress of the now extra-erythrocytic merozoites, which attach to and invade new RBCs to establish the next intra-erythrocytic proliferation cycle. This stage of the parasite life cycle is responsible for the disease pathology, and therefore is a principal target for the development of drugs to kill the parasite.

Several parasite proteins synthesized during this cycle are modified by *N*-myristoyl transferase (NMT). This enzyme transfers the C14 fatty acid from myristoyl-CoA to the amino terminal glycine of substrate proteins, in a largely co-translational event following removal of the initiator methionine (Dian, Perez-Dorado et al., 2020). Substrate proteins have been predicted bioinformatically, using a partially conserved sequence recognition motif (Castrec, Dian et al., 2018), or identified experimentally by metabolic incorporation of YnMyr, an alkyne-containing myristate analogue, which provides a convenient means to label such proteins and allow their purification and identification following the chemical addition of a biorthogonal tag (Broncel, Dominicus et al., 2020, Broncel, Serwa et al., 2015, Thinon, Serwa et al., 2014, Wright, Clough et al., 2014). Thirty-two *N*-myristoylated parasite proteins have been identified experimentally in the *P. falciparum* asexual blood stages (reviewed in (Schlott, Holder et al., 2018)). These NMT substrates are targeted to membranous structures such as the PM and the secretory pathway, which has a key role not only in protein export but also in the biogenesis and function of the IMC and intracellular organelles as well as protein import into the apicoplast (Schlott et al., 2018). Other myristoylated proteins are targeted to the nucleus or exported to the host erythrocyte. They function in a diverse range of cellular pathways such as protein secretion, transport and homeostasis, ion channel regulation and parasite motility, with their known enzymatic functions including kinase, phosphatase and hydrolase activities (Schlott et al., 2018). About one third of the experimentally identified NMT substrates were shown to be essential in parasite growth screens using insertional mutagenesis in *P. falciparum* (Zhang, Wang et al., 2018) and gene knockout in *Plasmodium berghei* (Gomes, Bushell et al., 2015, Schwach, Bushell et al., 2015), however this genetic evidence fails to indicate whether or not *N*-myristoylation is essential for the proteins’ function.

NMT inhibitors have been developed that kill the parasite *in vitro* (Bell, Mills et al., 2012, Rackham, Brannigan et al., 2014, Yu, Brannigan et al., 2012). Each of these inhibitor classes has been shown to bind to the protein substrate binding site of NMT, and their mode of action was confirmed using a parasite expressing a variant NMT with an amino acid substitution that abolishes both inhibition of enzyme activity and inhibition of parasite growth (Schlott, Mayclin et al., 2019). One such inhibitor, IMP-1002, when added to a synchronous population of ring stage parasites, interrupts parasite development irreversibly during early schizont development (four to six nuclei) and before the formation of the IMC, producing a parasite form that we have termed a ‘pseudoschizont’ (Wright et al., 2014). It is likely that this form results from inhibition of NMT in the trophozoite or early schizont stages. However, many NMT substrates are expressed abundantly later in schizogony and the consequence of NMT inhibition during this later stage, during a period of approximately 10 - 15 hours following commencement of nuclear division, is unknown. Protein myristoylation may result in increased membrane binding affinity; therefore, loss of myristoylation can cause aberrant subcellular targeting and consequent loss of protein function, and even degradation (Timms, Zhang et al., 2019). At the cellular level, inhibition of schizont development, for example through impaired nuclear division or defective formation of intracellular organelles, may prevent merozoite formation, parasite egress, merozoite invasion and subsequent ring stage development. To investigate these potential phenotypes, we examined the effect of inhibitor added during schizogony on parasite development, egress and invasion, and on myristoylated protein location and stability. The results showed that NMT inhibitor (NMTi) treatment during schizogony did not stop nuclear division, but it did inhibit parasite development before egress. We then developed a genetic method to examine whether the N-terminal glycine (and hence myristoylation) of a selected set of six proteins was essential for parasite growth. For members of the chosen panel of NMT substrates, substitution of N-terminal glycine with alanine was detrimental to parasite growth. From this set of proteins, we focused on one, glideosome-associated protein 45 (GAP45), to examine the importance of *N*-myristoylation for its localization and function. Induced replacement of the N-terminal glycine of GAP45 with alanine, had no effect on protein targeting to the IMC, the protein’s palmitoylation, or egress, but it did prevent merozoite invasion. We conclude that protein myristoylation is important at different time periods for nuclear division, merozoite maturation prior to egress, and for RBC invasion, implying that NMT inhibitors impact multiple facets of parasite development and are therefore excellent leads for drug development.

## RESULTS

To investigate the effect of an NMTi during schizogony, synchronized parasite populations were treated with either 140 nM IMP-1002 (the EC_90_ of the compound (Schlott et al., 2019)) or DMSO during the period from 34 to 45 h post invasion (PI), after which the culture medium was exchanged to a drug-free medium at the first sign of parasite egress in the DMSO-treated culture. Parasite growth, invasiveness and morphology were then assessed by flow cytometry of Hoechst-stained parasites and microscopy of Giemsa- or antibody-stained fixed parasites, while parasite proteomics were analysed by mass spectrometry.

### Parasite proliferation is decreased significantly by inhibition of NMT during schizont differentiation, blocking parasite development before egress

Flow cytometry analysis of samples stained with Hoechst dye demonstrated a significant drop in parasite proliferation resulting from IMP-1002 NMT inhibition compared with DMSO controls (p < 0.0001, Supplementary Figure 1). Therefore, both ring and schizont populations were examined separately, to determine whether this decrease resulted from reduced parasite egress from infected erythrocytes or defective invasion into new erythrocytes. Using Percoll-purified schizonts, two samples, eight hours (53 h PI) and twenty-eight hours (73 h PI) after the start of egress of control (DMSO-treated) parasites, were used to determine the growth rate (Figure 1A) and measure the schizont and ring stage parasitemia in each sample (Figure 1B). The growth rate dropped significantly in the presence of IMP-1002 compared with DMSO controls (p < 0.0001 for 53 h PI and p < 0.0003 for 73 h PI). In the DMSO-treated control culture the schizont population decreased and the ring population increased during the period between 45 and 73 h PI, indicating merozoite egress and invasion, whereas the schizont population in IMP-1002 treated samples remained constant and few ring stage parasites were detected by 73 h PI. These data indicate that NMT inhibition blocks parasite development in the schizont stage, before merozoite egress.

**Figure 1.**
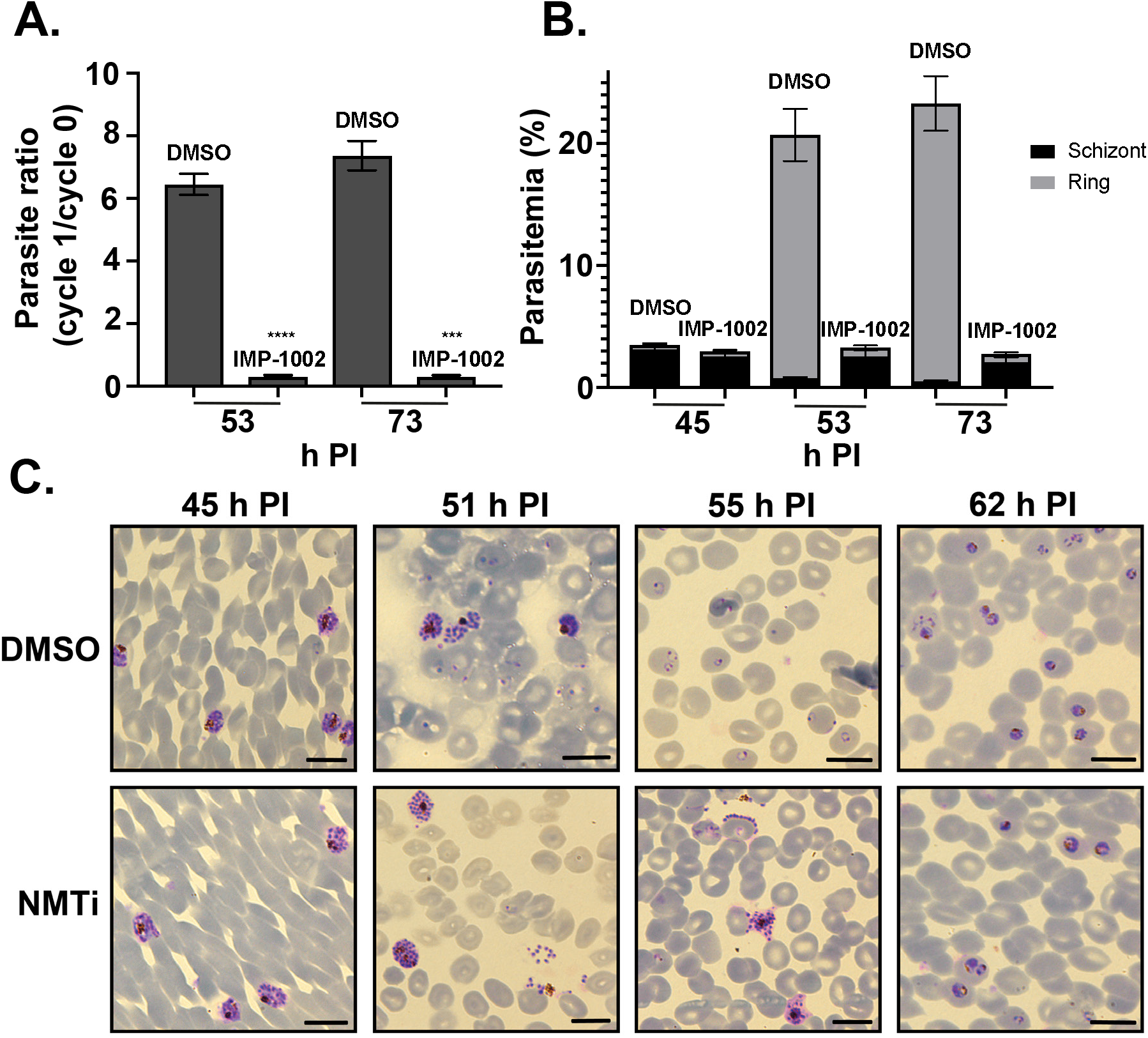
Inhibition of NMT during schizogony leads to a block in parasite development before merozoite egress from infected erythrocytes. **A.** The parasite ratio (parasitemia of cycle 1/ parasitemia of cycle 0) at 53 and 73 hours post infection (h PI) measured by flow cytometry. Equal numbers of IMP-1002- and DMSO-treated parasites, collected at 45 h PI and Percoll purified, were mixed with fresh erythrocytes for a growth assay. The growth of IMP-1002-treated parasites was significantly lower (p < 0.0001 for 53 h PI and p < 0.0003 for 73 h PI: unpaired Student t-test with Welch’s correction not assuming an equal SD, n = 3; 50.000 RBC counted per sample). **B.** Percentage of schizonts and rings in the growth assay samples. While the number of schizonts remained the same during the period from 45 h PI to 73 h PI and few rings were detected even at 73 h PI in the IMP-1002 inhibitor treated samples, the size of the schizont population dropped and the ring population increased substantially in the DMSO-treated control culture (n=3). **C.** Giemsa staining to reveal parasite morphology after eleven-hour drug treatment. While there was no visible morphological difference between IMP-1002- and DMSO-treated schizonts at 45 and 51 h PI, by 55 h PI abnormalities in schizont morphology became apparent in the drug-treated culture. There is an abnormal distribution of merozoites around the hemozoin in combination with a less spherical structure of the PM/PVM. Parasites that survived drug treatment developed normally into trophozoites (shown at 62 h PI) (20 fields of view per sample, n = 3),. Scale bar = 10μm.

Giemsa staining and microscopy indicated that at 45 and 51 h PI drug-treated schizonts looked similar morphologically to the DMSO-treated control parasites (Figure 1C). But at about ten hours after the exchange to a drug-free medium (at 55 h PI) IMP-1002-treated schizonts started to appear abnormal and there was little evidence of invasion and new ring stage formation, suggesting that these parasites were not viable (Figure 1C). However, at 62 h PI some parasites that had escaped IMP-1002 NMT inhibition had developed into healthy-looking trophozoites. These results, obtained by microscopy, complement those from the flow cytometry analysis and indicate that at IMP-1002 EC_90_, NMT inhibition blocks schizont development before merozoite egress in all but a small fraction of parasites.

### NMT inhibition changes substrate protein solubility and localization and disrupts rhoptry formation

NMT substrates may associate differently with membranes when parasites have been treated with NMTi. To examine this, schizonts were subjected to sequential solubility fractionation and the distribution of specific proteins was revealed by western blotting. Both armadillo domain-containing rhoptry protein (ARO) and calcium dependent protein kinase 1 (CDPK1), proteins that have an N-terminal myristoylation site and an adjacent potential palmitoylation site, were largely in the membrane-bound fraction prepared from DMSO-treated parasites (Figure 2A). However, following parasite treatment with IMP-1002, the proteins were either completely (in the case of ARO) or partially (in the case of CDPK1) found in the hypotonic buffer-soluble fraction. The IMC protein, GAP45, which has an N-terminal myristoylation site, an adjacent palmitoylation site, and an additional palmitoylation site near the C-terminus (Jones, Collins et al., 2012), showed no difference in its solubility profile following IMP-1002 NMT inhibition. As controls, we identified the fractions enriched for cytoplasmic heat shock protein 70 (HSP70) and myosin tail interacting protein (MTIP), a component of the glideosome, formed together with GAP45 and other proteins. As expected, HSP70 was largely in the soluble fraction and MTIP was in the membrane bound fraction, and their behaviour was not affected by IMP-1002 NMTi treatment of the parasite.

To examine the effect of NMT inhibition on the subcellular protein location during schizont development, parasites were analysed using an indirect immunofluorescence assay (IFA) with specific antibodies (Figure 2B to 2D). Following NMTi treatment, the location of ARO and rhoptry neck protein 4 (RON4), changed from being discrete to very diffuse in the cytoplasm of developing merozoites (Figure 2B). The IMC proteins, GAP45 and myosin A (MyoA) showed no discernible difference in their location and were present in both DMSO and IMP-1002 treated cells (Figure 2C). The subcellular location of the micronemal protein, erythrocyte binding antigen 175 (EBA-175), was also unaffected by IMP-1002 treatment, when compared to the DMSO control (Figure 2D).

**Figure 2.**
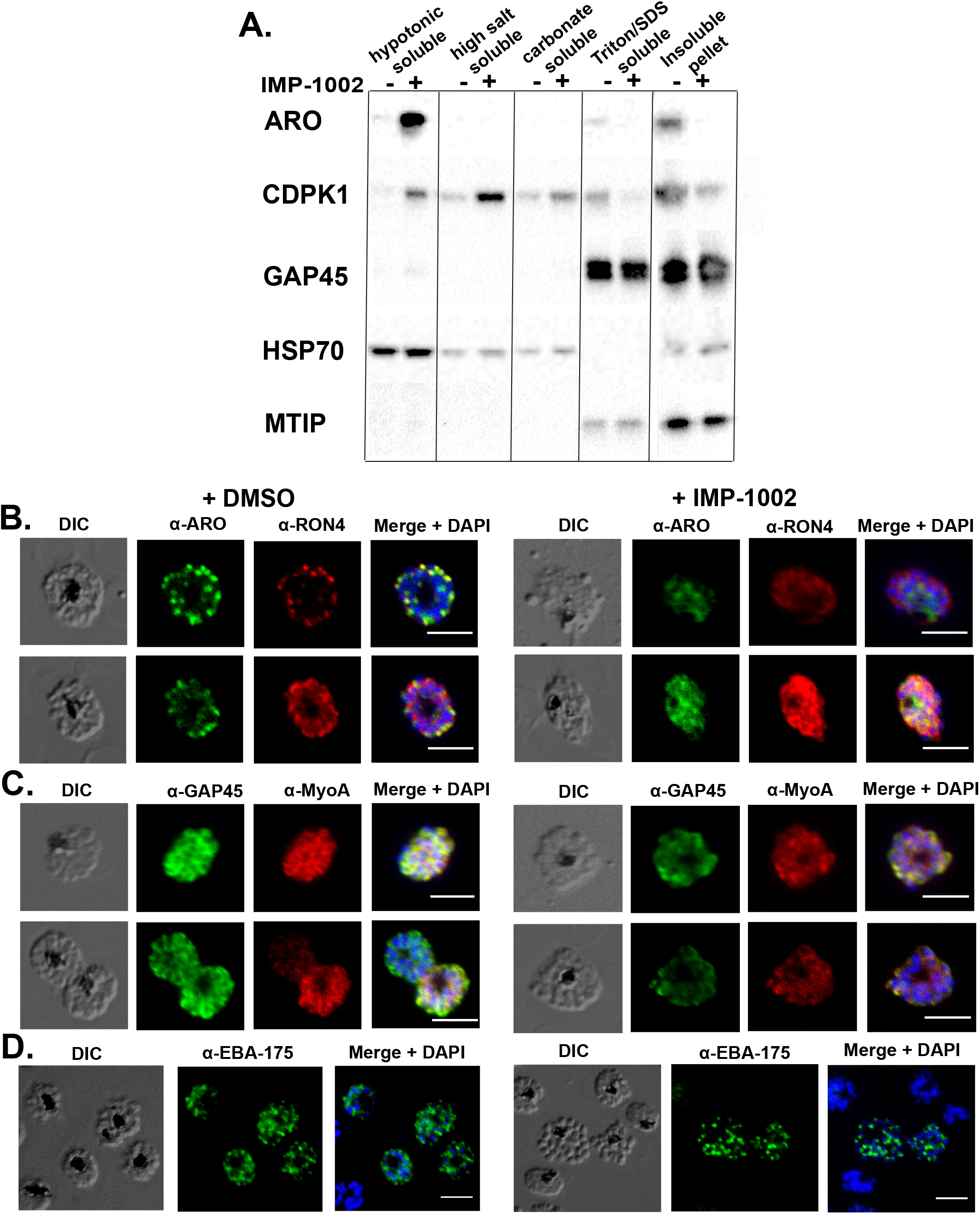
IMP-1002 treatment during schizogony changes the differential solubility of ARO and CDPK1 and the subcellular location of ARO and RON4. **A.** To examine the differential solubility of proteins present in IMP-1002-treated and untreated parasites, Percoll-purified schizonts were lysed and sequentially fractionated using hypotonic and high salt buffers, sodium carbonate, and a buffer containing 1% Triton X100 and 0.1% SDS. These fractions together with the insoluble pellet, were analysed by western blot using antibodies to ARO, CDPK1, GAP45, HSP70, MSP7 and MTIP. In the presence or absence of IMP-1002 ARO was largely in the hypotonic soluble and carbonate insoluble fractions, respectively; CDPK1 was distributed in the hypotonic/high salt soluble and carbonate insoluble fractions, respectively, under the same conditions. Microscopy images are from indirect immunofluorescence assays (IFAs) performed in duplicate on three separate occasions with fixed parasites from DMSO (control) and IMP-1002-treated parasites, and protein-specific antibodies. Panels show the differential interference contrast (DIC) image, the specific antibody location (green or red) and a merged image of the antibody staining with DAPI staining of nuclei. Scale bar = 5 μm. **B.** ARO and RON4 localization was affected by drug treatment, with the proteins distributed throughout the cytoplasm of developing merozoites and loss of distinct rhoptry staining. **C.** The location of GAP45 and MyoA at the inner membrane complex (IMC) appeared largely unchanged. **D.** The location of the micronemal protein EBA-175 was not affected by IMP-1002 treatment.

These results indicate that schizont treatment with IMP-1002 can affect the membrane binding properties of some proteins and, for example in the case of ARO, may result in their mislocalization within the cell.

### IMP-1002 inhibits protein myristoylation and affects abundance of some NMT-substrates and non-myristoylated proteins

While the localization of NMT substrates can be studied by cellular fractionation or microscopy-based approaches, these methods provide no quantitative data on the effect of IMP-1002 inhibition on the modification or abundance of parasite proteins. Therefore, we used quantitative chemical proteomics to examine further the effect of NMT inhibition on both myristoylation of its substrates and the abundance of other proteins in the cell. The extent of myristoylation was studied using metabolic labelling with the myristic acid analogue YnMyr and label-free quantification (LFQ) by mass spectrometry to determine the relative abundance of individual myristoylated proteins in samples from parasites treated with either DMSO or IMP-1002 for eleven hours. Proteins were extracted and an AzTB biotin tag attached to the YnMyr-labelled proteins using click chemistry, then the tagged proteins were enriched by Neutravidin binding and elution. A total of 609 proteins were identified in the eluate from the Neutravidin-coated agarose beads (Supplementary Data 1). Sixteen NMT substrates showing a significant decrease in myristoylation in the presence of IMP-1002 were identified (Figure 3A, Supplementary Data 1). Fourteen of these sixteen proteins were experimentally verified NMT substrates (Wright et al., 2014), while the two remaining proteins were a putative kinase (PF3D7_0321400) and a conserved protein of unknown function (PF3D7_0619700) (Figure 4A). The modified N-terminal glycine was also identified for a number of NMT substrates, providing direct experimental evidence of myristoylation, for example metal-dependent protein phosphatase 6 (PF3D7_1309200) and putative acylated pleckstrin-homology domain-containing protein (PF3D7_0414600) (Supplementary Figure 2). Glycosyl phosphatidylinositol (GPI) anchored proteins, which incorporate YnMyr through an ester linkage, and non-myristoylated IMC proteins were largely unchanged (analysis using adjusted p-values with an FDR of 0.01 and within group variance S_0_ = 0.5, n = 3).

**Figure 3.**
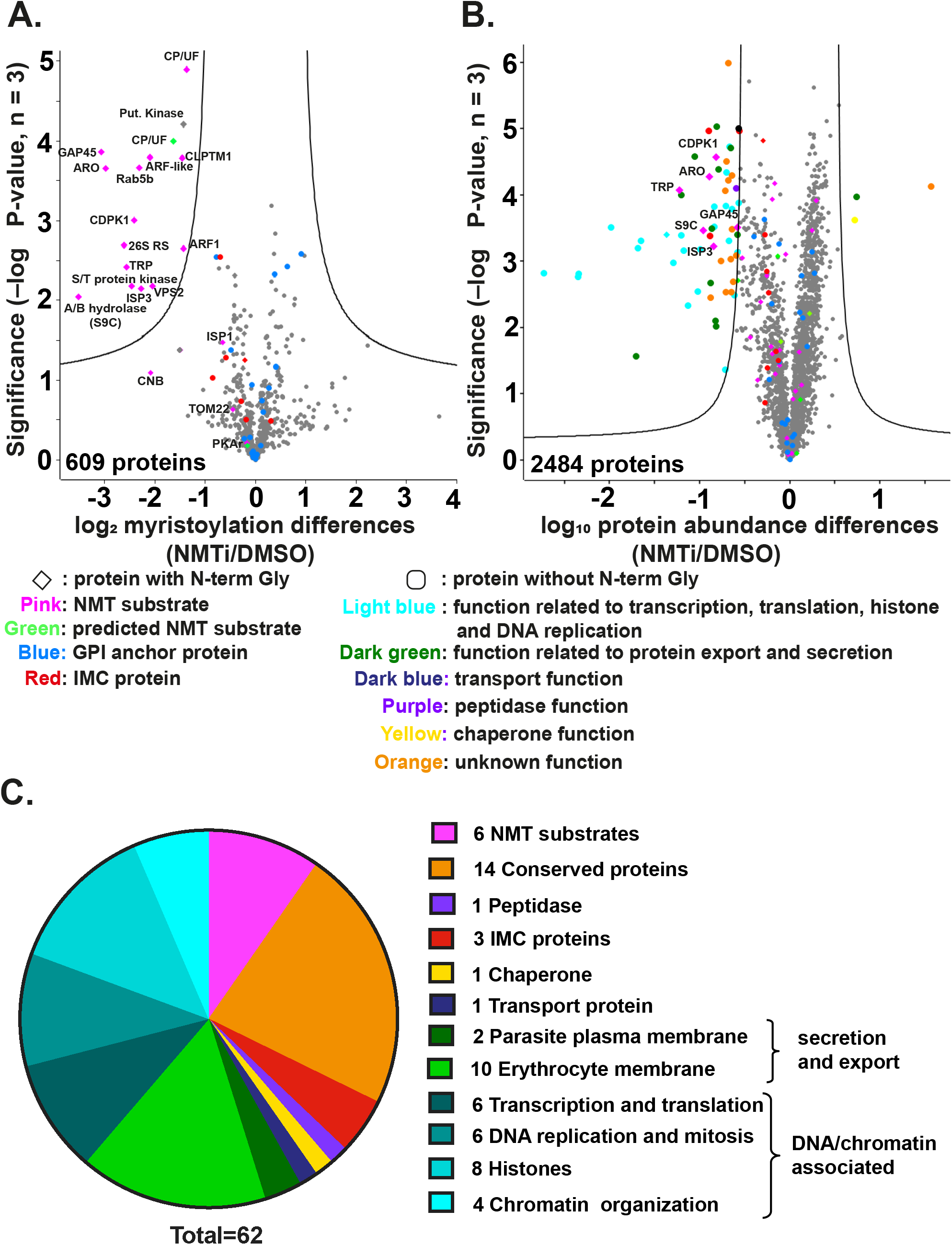
NMT inhibition changes the abundance of both myristoylated and non-myristoylated proteins. **A.** Parasite proteins were metabolically labelled with YnMyr for eleven hours during schizogony, coupled to AzTB and enriched on Neutravidin coated agarose beads. Label-free quantification (LFQ) analysis was used to measure the abundance of enriched proteins labelled in the presence or absence of IMP-1002. A two-sample t-test (permutation-based false discovery rate [FDR], 250 permutations [number of randomizations], FDR 0.01, S0 = 0.5 [within groups variance]; n = 3 biological replicates, each with three technical replicates) revealed significant differences in myristoylated protein abundance between IMP-1002-treated and control (DMSO) samples. The lines on the graph indicate t-test significance cut off. The identity of some proteins is shown on the plot; symbols and colour coding of individual proteins are explained below the plot. Full data are in Supplementary Data 1. **B.** Quantitative whole proteome analysis using tandem mass tag (TMT) protein labelling to measure protein abundance in parasite samples treated with either DMSO or 140 nM IMP-1002 and subsequent saponin lysis. A two-sample t-test (permutation-based FDR, 250 permutations, FDR 0.01, S0 = 0.8 [within groups variance], n = 3) revealed significant changes in overall protein abundance between the inhibitor-treated and control (DMSO-treated) parasites. The identity of some proteins is shown on the plot; symbols and colour coding of individual proteins are explained below the plot. Full data are in Supplementary Data 2. **C.** Pie chart presentation of the 62 proteins significantly reduced in abundance following IMP-1002 treatment of schizonts from 34 to 45 h PI, with their associated grouping based on GO term analysis from PlasmoDB (Release 44, July 2019).

**Figure 4.**
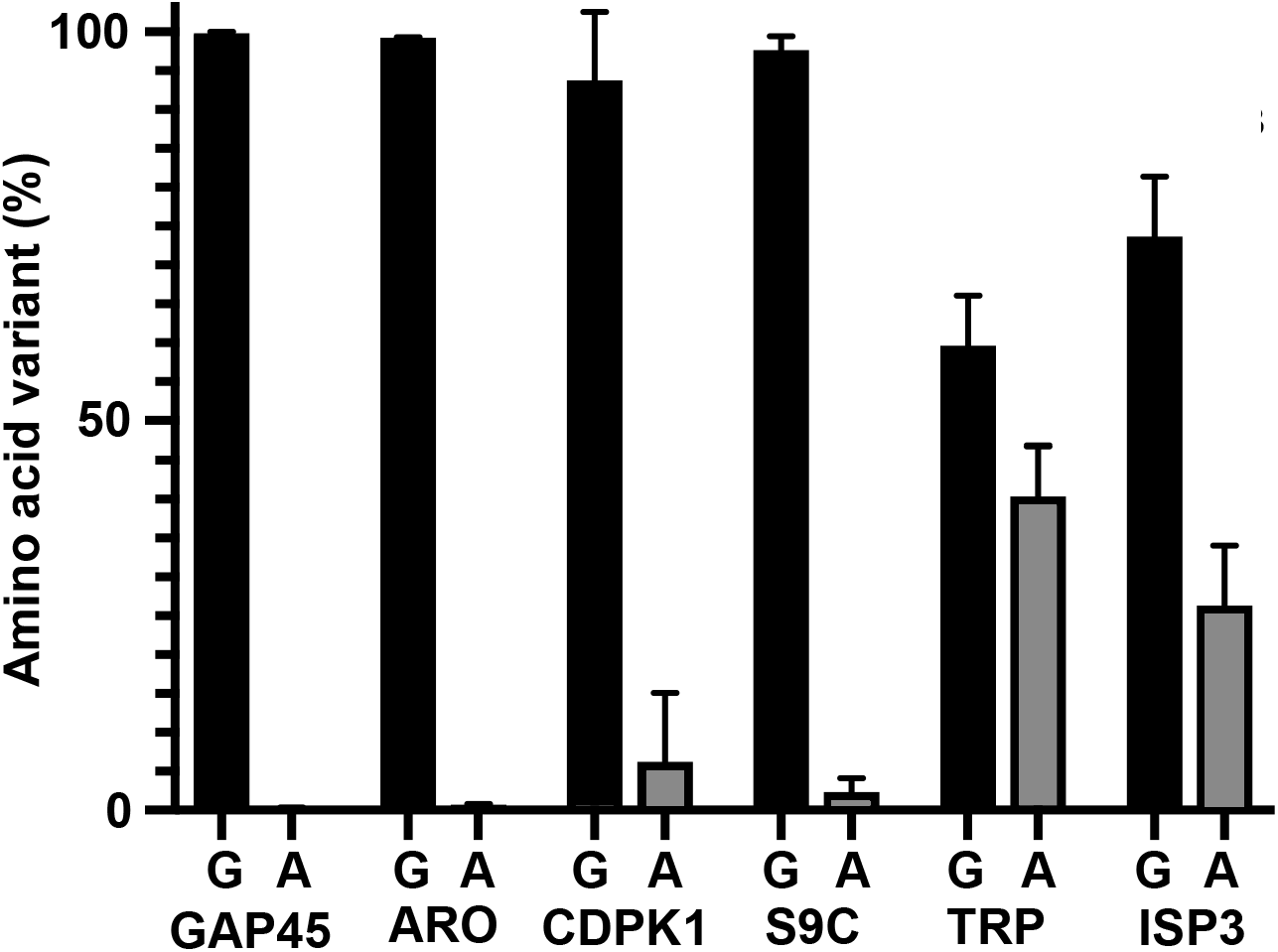
A viability screen of parasites containing either G2G or G2A at the myristoylation site in NMT substrates. **A** CRISPR-Cas9 approach was used to insert G2A or G2G codons at the start of selected genes (see Supplementary Figure 3 for details). Six genes were targeted using a 50:50 ratio of repair plasmids carrying a codon for G2G (Gly; silent mutation) or G2A (Ala, abolishing the myristoylation site). Illumina sequencing of products amplified using integration specific PCR primers was used to determine the distribution of each form in the parasite population following transfection. Each gene targeting was performed at least twice (*gap45*, n = 3) with two different guides (except for *aro* and *gap45* when two transfections with the same guide were performed).

To determine if there was an effect of NMT inhibition on overall protein abundance, the proteome of schizonts that had been incubated with or without 140 nM IMP-1002 from 34 – 48 h PI was analysed by mass spectrometry using tandem mass tag (TMT) labelling as a quantitative method, in combination with an additional high pH reverse fractionation step to increase coverage of the multiplex sample. A total of 2,484 proteins was identified (Supplementary Data 2), of which 62 were significantly reduced in abundance by IMP-1002 treatment (t-test using a false discovery rate (FDR) cut-off of 0.01 and a within-group variance (S0) of 0.8) (Figure 3B). Gene ontology (GO)-term analysis of these 62 proteins revealed six NMT substrates, together with several proteins involved in DNA replication and chromatin function, as well as a number of exported/secreted proteins (Figure 3C). The NMT substrates were ARO [GeneID: PF3D7_0414900], CDPK1 [PF3D7_0217500], GAP45 [PF3D7_1222700], alpha/beta hydrolase S9C [PF3D7_0403800], IMC sub-compartment protein 3 [ISP3; PF3D7_1460600] and tetratricopeptide-repeat proteins [TRP; PF3D7_0601600, PF3D7_0631000]. In previous studies, three of these proteins had been either suggested (CDPK1, ARO) (Zhang et al., 2018) or shown (GAP45 (Perrin, Collins et al., 2018)) to be essential for growth of *P. falciparum* asexual blood stage parasites. Insertional mutagenesis of alpha/beta hydrolase S9C produces a slow growing phenotype (Zhang et al., 2018) and *P. berghei* ISP3 is dispensable (Poulin, Patzewitz et al., 2013). Of the TRP-encoding genes, PF3D7_0631000 was classified as essential and PF3D7_0601600 was classified as dispensable in a recent *P. falciparum* mutagenesis screen, (Zhang et al., 2018). However, none of these earlier studies addressed the essentiality of the N-myristoylation.

These data show that IMP-1002 treatment has a direct effect on the myristoylation of NMT substrates. Furthermore, there is a reduced abundance of both myristoylated and non-myristoylated proteins in the treated cells compared to those incubated with DMSO. There were six myristoylated proteins that were significantly reduced in abundance, suggesting that these NMT substrates are of particular importance. Therefore, we developed a genetic screen to look specifically at the importance of the N-terminal glycine of these proteins, and hence myristoylation, on parasite growth.

### A G2A/G2G CRISPR-Cas9 screen identifies substrates for which myristoylation is required for parasite viability

The six NMT substrates significantly reduced in abundance by IMP-1002 treatment during schizogony were selected for further analysis to examine whether or not N-terminal myristoylation is essential for parasite growth. We developed a CRISPR-Cas9 screen to determine the relative fitness of parasites following integration of a G2A codon or a G2G replacement codon at the myristoylation site for each of the six substrates. For TRP we used PF3D7_0631000 for this screen, as it has been shown to be essential for parasite viability (Zhang et al., 2018). For each gene, Cas9 was used to generate a double-strand break within the coding region close to the 5’ end of the gene using two different guides (Supplementary Figure 3, Supplementary Table 1), with repair mediated by plasmids containing either a G2A sequence to prevent myristoylation of the protein or a G2G sequence to allow it. Repair plasmids were mixed in equal proportion and added together to Cas9/guide plasmids, linearized and used for parasite transfection. Then at the same time post-transfection, parasite genomic DNA was extracted and integration-selective primers annealing to the inserted recodonized repair sequence were used to attach adapter sequences for Illumina sequencing (Supplementary Table 2). The ratio of G2A/G2G sequence reads for each parasite culture provides an indication of the relative viability of the G2G and G2A variants; with no fitness cost a 1:1 ratio of the two forms would be expected in the parasite population.

For four of the six genes, almost 100% of the retrieved integrated sequences coded for an N-terminal glycine, suggesting that the N-terminal glycine is essential for these proteins (Figure 4), although the number of reads recovered for the CDPK1 gene was small (Supplementary Figure 3). For the TRP gene only 60% of the reads were for the N-terminal glycine sequence and for ISP3, a 26% incorporation of the G2A variant was detected. Overall, the screen showed that for four out of the six NMT substrates, myristoylation is likely essential for viability (GAP45, ARO, CDPK1, and S9C), while for two substrates (ISP3 and TRP) myristoylation might be dispensable. To carry this analysis further we focused on one protein, GAP45, which has been shown to be essential for motor complex formation and invasion, and used a genetics-based complementation approach to investigate the importance of GAP45 N-myristoylation.

### The N-terminal glycine of GAP45 is essential for parasite viability

We focused on GAP45, for which myristoylation appears to be essential for viability, for further detailed analysis of the consequence of lack of myristoylation. The strategy used a genetic complementation approach by further modification of an existing parasite that had been engineered to allow an inducible knockout of *gap45*. The *gap45* gene has been shown to be essential in the *gap45:ha3:loxP* parasite line, which has a loxPint intron after the first 49 base pairs, a loxP site after the stop codon, expresses HA-tagged GAP45 and allows an inducible knockout of the gene (Perrin et al., 2018). We inserted a second copy of the *gap45* gene together with its own promoter sequence into the *pfs47* locus to express either a wild type (WT) GAP45 or GAP45[G2A] gene, and examined whether or not this second gene complemented the induced knockout of *gap45*. Construction of these parasite lines is shown in Supplementary Figure 4. For both transfections, parasites were detected after 22 days and following confirmation of DNA integration, parasite lines were cloned by limiting dilution. The parasite clones used for complementation analysis are denoted as *gap45:ha3:loxP∷comp_gap45[WT]* and *gap45:ha3:loxP∷comp_gap45[G2A]*, respectively.

First we examined protein levels of the *gap45:ha3:loxP*, *gap45:ha3:loxP∷comp_gap45[WT]* and *gap45:ha3:loxP∷comp_gap45[G2A]* schizonts by western blotting and IFA using anti-GAP45 antibodies. The HA-tagged GAP45 is 4.3 kDa larger than GAP45 expressed at the same time from the second gene copy, which has no HA-tag (Perrin et al., 2018), and therefore both forms were visible on a western blot with anti-GAP45 antibodies (Figure 5A). In cycle 0, the cycle in which rapamycin treatment was given to induce gene excision (Supplementary Figure 5), expression of GAP45 in the *gap45:ha3:loxP* line was undetectable, whereas in the *gap45:ha3:loxP∷comp_gap45[WT]* and *gap45:ha3:loxP∷comp_gap45[G2A]* lines GAP45 was present at approximately the same levels as in the DMSO treated controls. The IFA analysis confirmed that rapamycin treatment abolished expression of the HA-tagged protein and that the gene inserted into the Pfs47 locus produces GAP45 that is indistinguishable in location from wild type GAP45 (Figure 5B). By morphology, comparing the *gap45:ha3:loxP, gap45:ha3:loxP∷comp_gap45[WT]* and *gap45:ha3:loxP∷comp_gap45[G2A]*, these lines developed normally through cycle 0, confirming the previous observation that full-length GAP45 is not essential for schizont development (Perrin et al., 2018). This result also demonstrates that loss of N-terminal myristoylation in GAP45 does not alter the subcellular localization of this modified protein.

**Figure 5.**
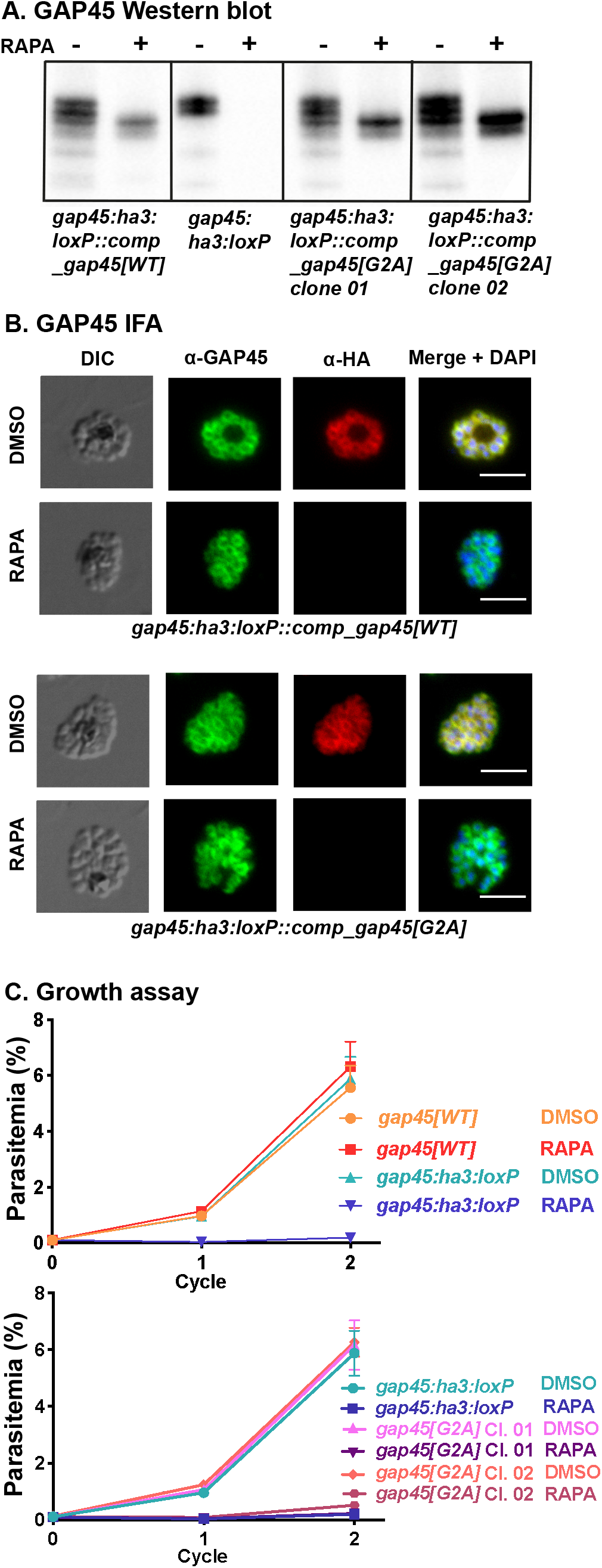
GAP45[G2A] is at the same subcellular location as GAP45, but the parasite has a growth defect. **A.** Western blots showing successful rapamycin-inducible ablation of *gap45:ha3:loxP* expression, but expression of wild type (WT) and G2A GAP45 from the Pfs47 site is unaffected. Note that HA3-tagged GAP45 adds additional mass to the protein resulting in a lower mobility in the gel. After rapamycin treatment, in GAP45[WT] and both GAP45[G2A] parasite clones the endogenous GAP45-HA3 is deleted but the second copy of the gene is still expressed (experiment performed in duplicate with two independent clones). **B.** In the presence of rapamycin, the endogenous HA-tagged GAP45 protein is no longer present but GAP45 expressed from the second gene copy in the Pfs47 locus is located at the periphery of the developing intracellular merozoites, as judged by IFA. In the presence of DMSO, GAP45-HA3 is expressed at this subcellular location as are GAP45[WT] and GAP45[G2A] in the presence or absence of rapamycin and DMSO for the *gap45:ha3:loxP∷comp_gap45[WT]* and *gap45:ha3:loxP∷comp_gap45[G2A]* parasite clones, respectively (experiment performed in duplicate with two independent clones). Scale bar, 5 μm. **C.** Growth of parasite lines following rapamycin or DMSO treatment over two cycles of development. Growth curves showing replication of the *gap45:ha3:loxP* parasite line following rapamycin or DMSO treatment. Rapamycin induced excision of the *gap45:ha3:loxP* locus produced parasites that were unable to replicate *in vitro* which can be complemented by the *gap45:ha3:loxP∷comp_gap45[WT]* but not by either of the two *gap45:ha3:loxP∷comp_gap45[G2A]* clones (gap45[G2A] Cl. 01 and Cl. 02). Means from three replicates plotted. Error bars show standard deviation.

In subsequent cycles after rapamycin treatment, however, GAP45 expressed in the *gap45:ha3:loxP∷comp_gap45[G2A]* integrant was not able to complement the *gap45:ha3:loxP* defect. After two cycles, neither rapamycin treated *gap45:ha3:loxP* nor *gap45:ha3:loxP∷comp_gap45[G2A]* parasites were able to proliferate (Figure 5C). In contrast, the rapamycin treated *gap45:ha3:loxP∷comp_gap45[WT]* parasites, and all three parasites treated with DMSO alone, continued to replicate. These data indicate that the N-terminal glycine and hence the myristoylation of GAP45 is indispensable for the survival of asexual blood-stage parasites.

### GAP45[G2A] assembles into an intact glideosome and is*S*-palmitoylated but not*N*-myristoylated

With these three parasite lines, we were able to investigate the role of GAP45 in glideosome assembly and its post-translational acylation. Previously, it had been shown that an N-terminal truncated GAP45, expressed together with WT GAP45, is incorporated into the glideosome (Ridzuan, Moon et al., 2012), but in the absence of GAP45 the glideosome does not form (Perrin et al., 2018). In the *Plasmodium* glideosome model, based on the *T. gondii* assembly, the C-terminal domain of GAP45 interacts with MyoA and its light chains, MTIP and ELC, and binds to the IMC via GAP50, playing a role in motility and host cell invasion (Frenal, Polonais et al., 2010). In the absence of GAP45, MTIP and MyoA are present at low levels and are not associated with the IMC (as assessed by IFA), although IMC formation is not impaired (Perrin et al., 2018), leading to the conclusion that GAP45 is essential for correct motor complex assembly, but not for maintaining the structural integrity of the IMC in *Plasmodium* (Perrin et al., 2018).

We examined the glideosome structure in *gap45:ha3:loxP, gap45:ha3:loxP∷comp_gap45[WT]* and *gap45:ha3:loxP∷comp_gap45[G2A]* parasites by IFA and western blot with or without rapamycin treatment. After rapamycin treatment of *gap45:ha3:loxP∷comp_gap45[G2A]* parasites, both MyoA and MTIP proteins were detected in the correct location by IFA (Figure 6A) and at normal levels by western blot (Figure 6B), in contrast to the situation in *gap45:ha3:loxP* parasites where MyoA and MTIP were not detectable. The location and abundance of GAP50 were unaffected. These findings indicate that although GAP45 is important for recruiting MyoA and MTIP to the IMC, its N-terminal glycine and hence its myristoylation is not required for this activity.

**Figure 6.**
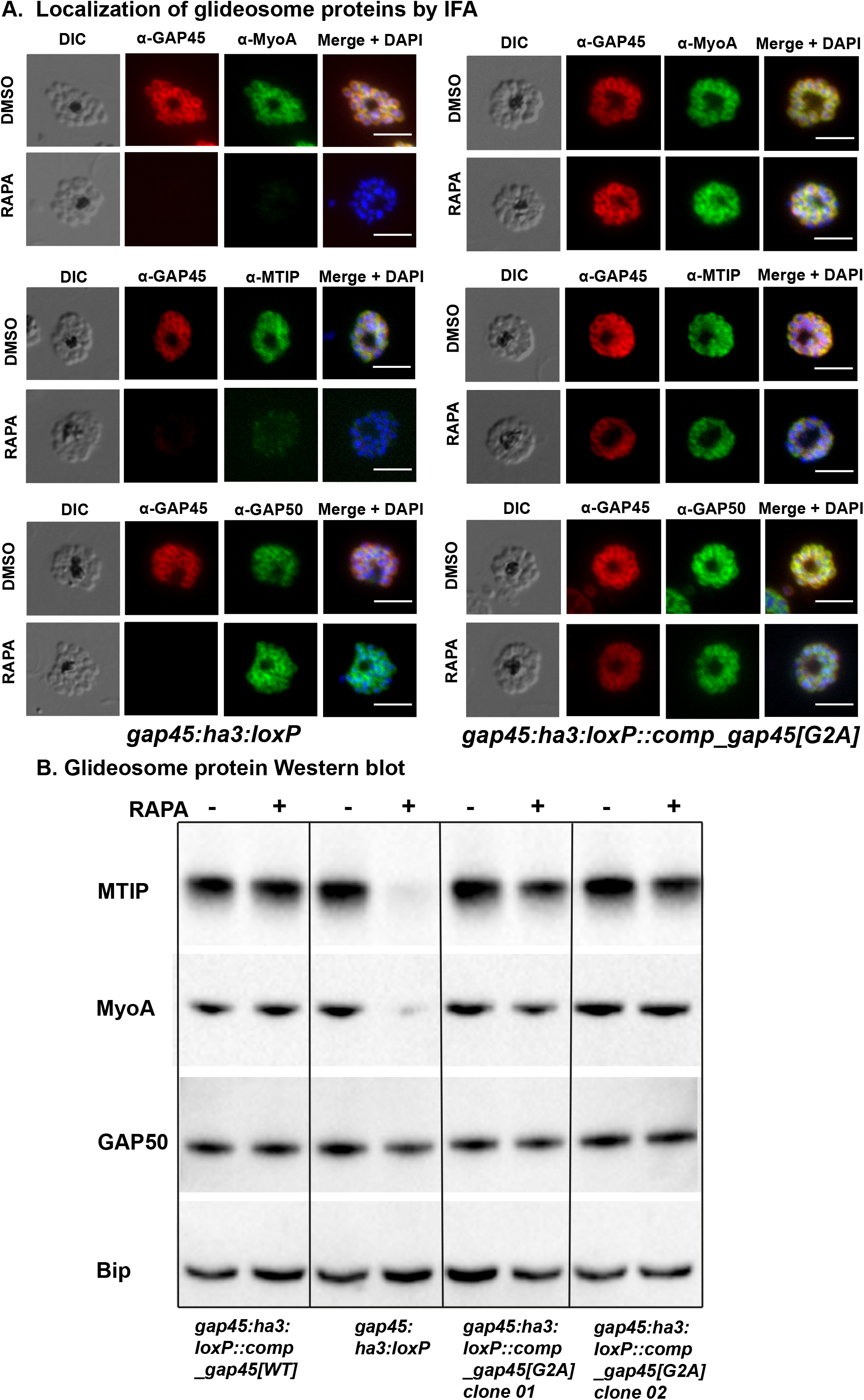
GAP45[G2A] parasites have a correctly localized glideosome. **A.** GAP45[G2A] parasites show no defects in expression of the glideosome components MyoA and MTIP, and which localized correctly at the periphery of merozoites. IFA showing the subcellular localization of GAP45-HA3, MyoA, MTIP and GAP50 in segmented schizonts of ∆GAP45 (*gap45:ha3:loxP*) and GAP45[G2A] (*gap45:ha3:loxP∷comp_gap45[G2A]*) in rapamycin and mock-treated (DMSO) parasites. Loss of GAP45 resulted in loss of detection of MTIP and MyoA at the IMC upon rapamycin treatment, while GAP45[G2A] is still able to recruit MTIP and MyoA. GAP50 staining is unchanged in both lines after rapamycin treatment (experiment performed in duplicate with two independent clones). Scale bars, 5 μm. **B.** Deletion of the GAP45 gene results in loss of the glideosome proteins, MTIP and MyoA in the *gap45:ha3:loxP* line after rapamycin treatment, but they are retained in the *gap45:ha3:loxP∷comp_gap45[WT] and gap45:ha3:loxP∷comp_gap45[G2A]* lines, as revealed by Western blotting. GAP50, another glideosome protein and the endoplasmic reticulum protein, Bip are unaffected by the GAP45 gene deletion. Experiment performed in duplicate with two independent clones.

To examine myristoylation, *gap45:ha3:loxP∷comp_gap45[G2A]* (2 clones)*, gap45:ha3:loxP* and *gap45:ha3:loxP∷comp_gap45[WT]* parasites were treated with rapamycin, or DMSO, during cycle 0, metabolically labelled with YnMyr from 34 h PI and harvested at 48 h PI. Proteins were extracted and an AzTB biotin tag attached to the YnMyr-labelled proteins using click chemistry. Tagged proteins were enriched by Neutravidin binding and then analysed by western blotting (Figure 7A). GAP45 was detected in the lysates and enriched protein fraction from all parasites treated with DMSO. However, after rapamycin treatment, only the *gap45:ha3:loxP∷comp_gap45[WT]* parasite expressed the myristoylated protein. As a positive control for myristoylation and the enrichment procedure, the known NMT substrate, ADP-ribosylation factor (ARF1, PF3D7_1020900), was successfully enriched, the additional mass of YnMyr conjugated to AzTB resulting in a small mobility shift, whereas, as a negative control the non-myristoylated endoplasmic reticulum chaperone BiP (PF3D7_0917900), was not enriched. These results indicate that, as predicted, GAP45[G2A] is not myristoylated.

**Figure 7.**
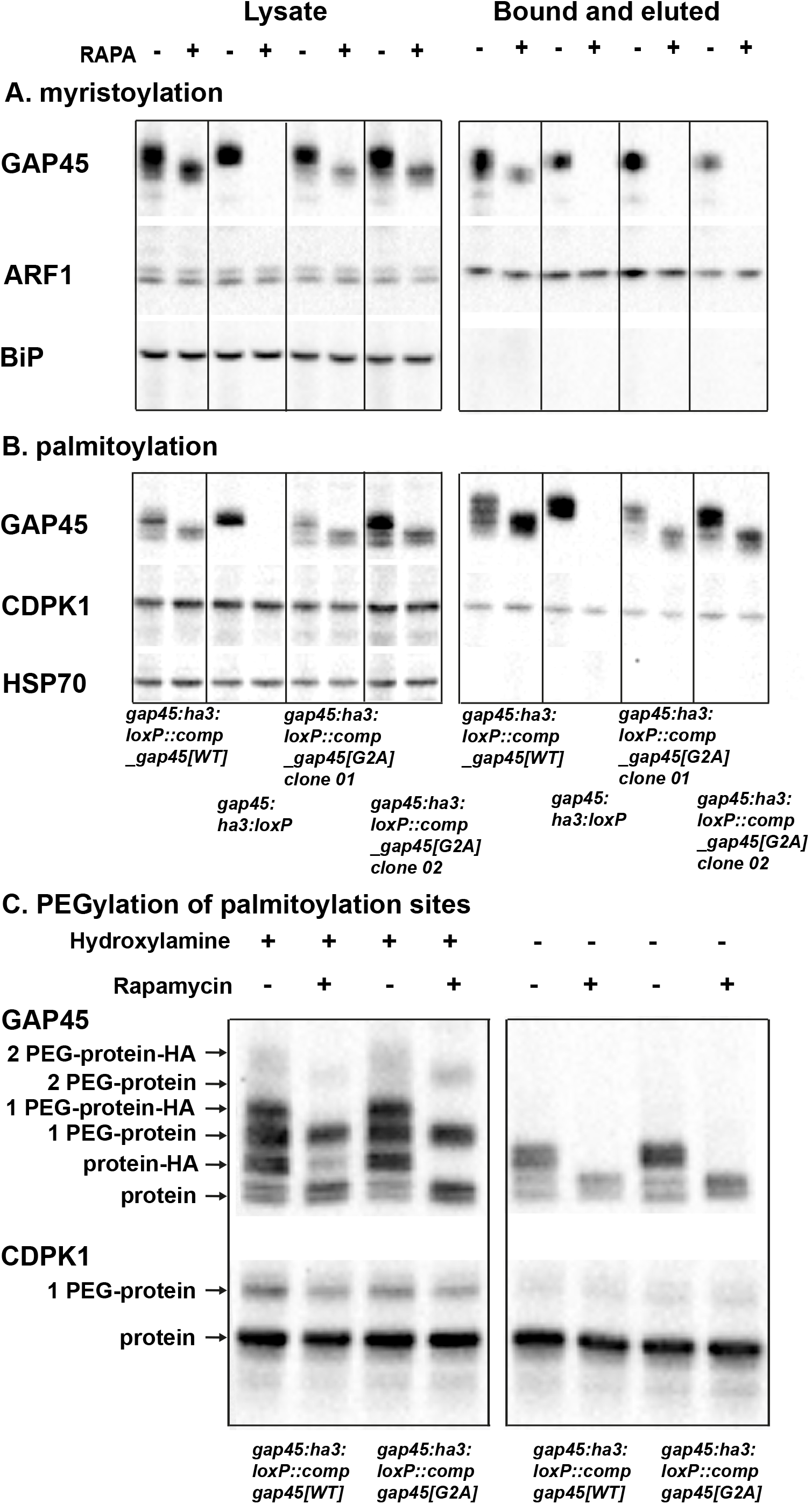
GAP45[G2A] is not myristoylated but is palmitoylated at a level similar to that of GAP45[WT]. Parasites were metabolically labelled in the presence or absence of rapamycin with either YnMyr (panel **A.**, myristoylation) or YnPal (Panel **B.**, palmitoylation), then the capture reagent AzTB was attached using CuAAC (click)-chemistry and the labelled proteins were enriched on Neutravidin. **A.** Western blot with anti-GAP45, anti-ARF1 and anti-BIP antibodies of enriched myristoylated proteins; a: lysate before enrichment, b: after enrichment: bound and eluted. **B.** Western blots with anti-GAP45, anti-CDPK1 and anti-HSP70 antibodies of enriched palmitoylated proteins; a: lysate before enrichment, b: after enrichment: bound and eluted. After rapamycin treatment only the *gap45:ha3:loxP∷comp_gap45[WT]* clone contained myristoylated GAP45. All other parasites showed no signal with the anti-GAP45 antibody after rapamycin treatment. The NMT substrate ARF1 was used as a positive control and was enriched from all four parasite clones, while the negative control BIP was not enriched as it is not myristoylated. All parasite clones except *gap45:ha3:loxP* expressed enriched palmitoylated GAP45 after rapamycin treatment, indicating palmitoylation of GAP45[G2A]. CDPK1 was used as a positive control and was enriched for all four clones, while HSP70 was not enriched as it is not palmitoylated. The experiment was carried out with two independent clones of gap45:ha3:loxP∷comp_gap45[G2A]. **C.** Acyl-PEG exchange (APE) reveals site-specific S-fatty acid acylation of GAP45[G2A] at a similar level to that of GAP45. *gap45:ha3:loxP∷comp_gap45[WT]* and *gap45:ha3:loxP∷comp_gap45[G2A]* parasites were treated with either rapamycin or DMSO and lysed at 48 h PI. Lysates were then subjected to APE, with or without hydroxylamine treatment to cleave esters, separated by SDS/PAGE, and analysed by Western blot with antibodies to either GAP45 or CDPK1. The mass of GAP45 is shifted by addition of the HA tag to the protein expressed from the endogenous locus, which is absent from the protein expressed from the *pfs47* locus. The number of PEGylation events is indicated. There was no evidence for a third palmitoylation of GAP45. CDPK1 was used a positive control; it has one palmitoylation site, visualised by the one PEGylation.

Since GAP45[G2A] is not myristoylated its modification by palmitoylation was examined. GAP45 has six cysteines that are potential sites for this modification: one at the N-terminus (Cys5), and five close to the C-terminus, of which one has been shown experimentally to be palmitoylated (Jones et al., 2012). The four parasite lines were synchronised, rapamycin treated and metabolically labelled with YnPal (heptadec-17-ynoic acid, also known as YnC14) to allow a biotin tag to be attached and the proteins enriched with Neutravidin coated beads. Samples were analysed by western blot using anti-GAP45, anti-CDPK1 (CDPK1 has a single palmitoylation site; a positive control), and anti-HSP70 (used as a negative control as HSP70 has no palmitoylation site) (Figure 7B). GAP45 was present in all samples except those from rapamycin treated *gap45:ha3:loxP* parasites. CDPK1 was present in all fractions including the enriched palmitoylated sample, whereas HSP70 was absent from the palmitoylated protein fraction. These results indicate that GAP45[G2A] is palmitoylated, to a similar extent as GAP45[WT], and that this modification is independent of prior myristoylation of the protein.

Incorporation of YnPal provides no indication of the number of palmitoylated cysteines in individual proteins. Therefore, to examine how many cysteines are palmitoylated in the different GAP45 proteins, we used acyl-PEG exchange (APE) methodology (Percher, Ramakrishnan et al., 2016). Proteins in cell extracts were reduced, reactive cysteine residues capped with *N*-ethylmaleimide (NEM), and then acyl thioester bonds were cleaved with hydroxylamine treatment, to allow site-specific alkylation with a 10 kDa methoxy(polyethylene glycol)-maleimide (mPEG-Mal) mass-tag. Each tag addition to a former palmitoylation site results in a discrete mobility shift detected on a western blot with anti-GAP45 antibodies (Figure 7C). The samples were split after NEM treatment and then either treated with hydroxylamine, or left untreated to reveal the background of mPEG-Mal tagging. The *gap45:ha3:loxP∷comp_gap45[G2A]* and *gap45:ha3:loxP∷comp_gap45[WT]* parasites were used, with and without rapamycin treatment. In the absence of rapamycin, HA-tagged GAP45, GAP45[G2A] and GAP45[WT] were all tagged with mPEG-Mal, and after rapamycin treatment only GAP45[G2A] and GAP45[WT], were labelled. The western blot suggested two 10 kDa band shifts, consistent with the addition of two mPEG-Mal moieties to both GAP45[WT] and GAP45[G2A], although the upper band was faint in both cases, and there was no evidence of a third site (Figure 7C). The intensity of each band quantified with ImageJ, confirmed that the single palmitoylation species was the most abundant modified form of the protein (Supplementary Table 3). CDPK1 was used as a control protein and displayed a single mPEG-Mal shift, consistent with a single palmitoylation site (Figure 7C, Supplementary Table 3).

### Myristoylation of GAP45 is dispensable for egress but essential for RBC invasion

The growth assay over two generations indicated that *gap45:ha3:loxP∷comp_gap45[G2A]* parasites had a severe growth defect (Figure 5), and previous work has shown that parasites lacking GAP45 are able to egress but not invade (Perrin et al., 2018). Therefore, the ability of *gap45:ha3:loxP∷comp_gap45[G2A]* parasites to egress and invade after rapamycin treatment was investigated. Giemsa-stained thin blood smears of rapamycin treated *gap45:ha3:loxP∷comp_gap45[G2A]* parasites at 48 h PI revealed increased numbers of free merozoites and a lack of ring-stage parasites when compared to the DMSO-treated control (Figure 8A), a pattern similar to that observed with *gap45:ha3:loxP* parasites. An invasion assay was used to compare *gap45:ha3:loxP∷comp_gap45[G2A], gap45:ha3:loxP∷comp_gap45[WT]* and *gap45:ha3:loxP* parasites, treated with either rapamycin or DMSO. *gap45:ha3:loxP∷comp_gap45[WT]* parasites were able to invade erythrocytes normally after rapamycin treatment, but *gap45:ha3:loxP∷comp_gap45[G2A]* and *gap45:ha3:loxP* showed significantly reduced invasion (parasitemia of cycle 1 / parasitemia of cycle 0; *p* = 0.001 and *p* < 0.0001, respectively; Welch’s unpaired two tailed t-test) (Figure 8B). In an assay with purified schizonts, schizont parasitemia had decreased after 24 h when compared to the parasitemia at 0 or 4 h, consistent with egress occurring normally (Figure 8C), but following rapamycin treatment both *gap45:ha3:loxP∷comp_gap45[G2A]* and *gap45:ha3:loxP* parasite cultures contained significantly fewer new ring stages after 24h in the first cycle (Welch’s unpaired two tailed t-test; *p* = 0.0042 for *gap45:ha3:loxP∷comp_gap45[G2A]* and *p* < 0.0002 for *gap45:ha3:loxP*). These data indicate that myristoylation of GAP45 is necessary to generate the functional GAP45 that is essential for successful erythrocyte invasion.

**Figure 8.**
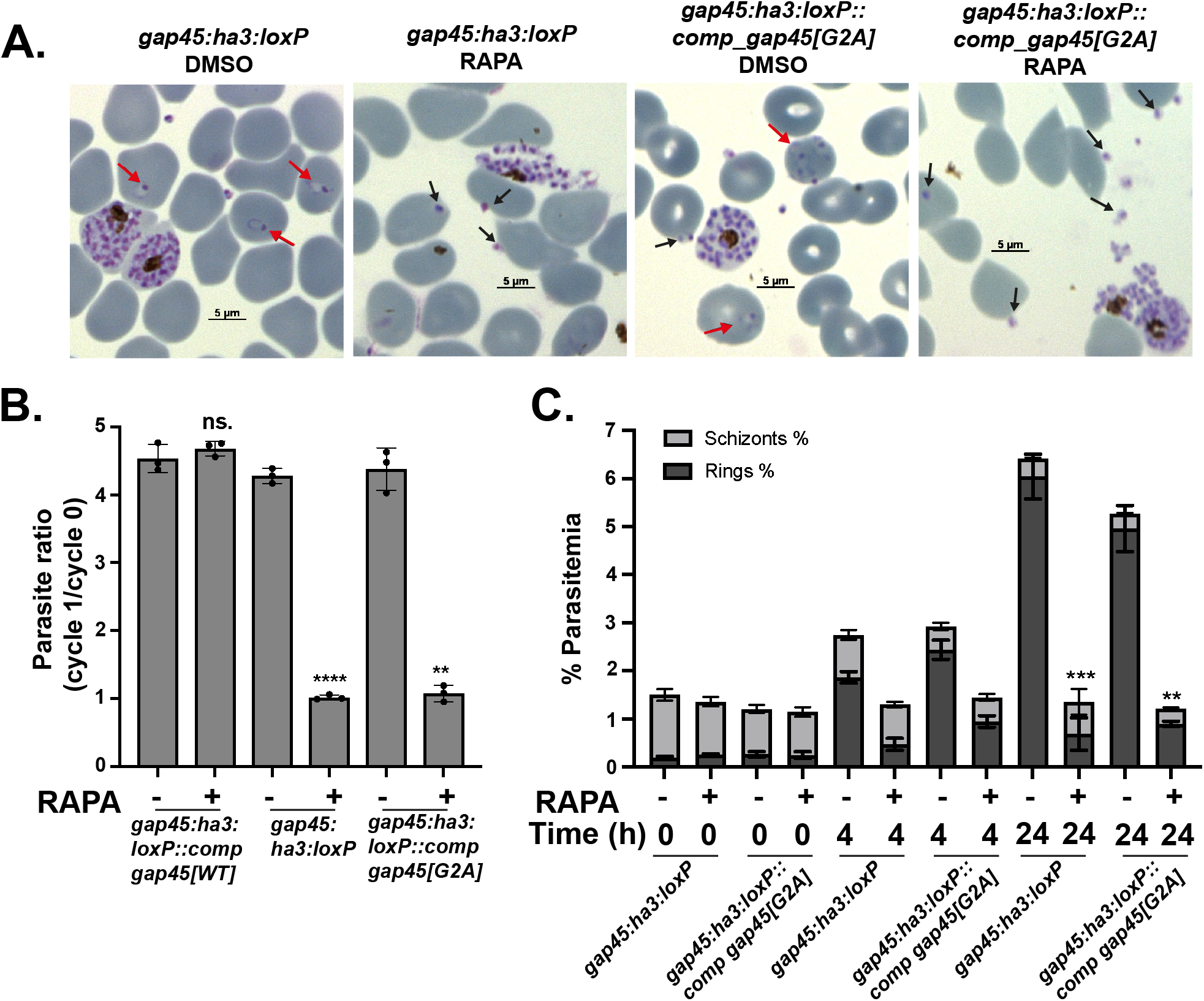
Parasites expressing GAP45[G2A] are not blocked at egress but show a defect in invasion. **A.** Giemsa-stained thin blood smears of *gap45:ha3:loxP* and *gap45:ha3:loxP∷comp_gap45[G2A]* clones treated with DMSO or rapamycin, showing the absence of newly invaded ring stages after rapamycin treatment in both lines, while invasion occurs in the DMSO control (red arrows). Despite no invasion after rapamycin treatment there is an abundance of extracellular merozoites visible with some apparently attached to erythrocytes (black arrows) indicating egress had occurred. **B.** Invasion assay of *gap45:ha3:loxP∷comp_gap45[WT]* (GAP45[WT])*, gap45:ha3:*loxP (∆GAP45), and *gap45:ha3:loxP∷comp_gap45[G2A]* (GAP45[G2A]), and showing the parasite ratio (parasitemia of cycle 1 / parasitemia of cycle 0) with and without rapamycin treatment (n = 3, Welch’s unpaired two tailed p < 0.0001 for KO and p = 0.001 for G2A); error bars show standard deviation. **C.** Percentage parasitemia of schizonts and rings in *gap45:ha3:*loxP (∆GAP45) and *gap45:ha3:loxP∷comp_gap45[G2A]* (GAP45[G2A) parasites at 0, 4, and 24 h after culture of schizonts grown in the presence of DMSO or rapamycin (n = 3 technical replicates with two independent clones, Welch’s unpaired two tailed t-test p < 0.0002 for KO and p = 0.0042 for G2A); error bars show standard deviation.

## Discussion

The consequences of NMT inhibition with IMP-1002 for *Plasmodium falciparum* depend on the length of incubation with the inhibitor and its concentration, as well as the stage of parasite development to which the inhibitor is added. Over 30 different NMT substrates have been identified experimentally in the asexual erythrocytic stage and just over 100 proteins of the total *Plasmodium* proteome are predicted to be myristoylated (Wright et al., 2014). Whilst many of the known NMT substrates likely have an essential function for the erythrocyte life cycle, the importance of *N*-myristoylation for that function is less clear, making it difficult to determine the key abnormalities resulting from NMT inhibition. One consequence of NMT inhibition for intra-erythrocytic stages was the failure to assemble the IMC during early schizogony, leading to a block in development. This is likely due to the lack of myristoylation of IMC components such as GAP45, ISP1 and ISP3 resulting in the formation of pseudoschizonts - cells with only four to five nuclei which fail to undergo further karyogenesis and cytokinesis (Wright et al., 2014). However, many proteins that are *N*-myristoylated are highly expressed during schizogony, in the last phase of intraerythrocytic development, and therefore we wished to examine in detail the consequence of treatment with NMTi specifically at this stage.

When IMP-1002 treatment was restricted to the last eleven hours of the cycle, parasites developed to schizonts that appeared morphologically fully mature, but parasite egress was prevented, leading to a significant drop in parasitemia in the subsequent cycle. Although there was no visible morphological change in parasites stained with Giemsa’s reagent, subcellular protein fractionation and immunofluorescence analysis showed that IMP-1002 treatment led to protein mislocalization. For example, the NMT substrate ARO is typically found in the membrane fraction, but in the presence of IMP-1002 it was soluble. A similar change in membrane association was displayed by CDPK1, but not by GAP45, which remained membrane bound, likely due to its additional *S*-palmitoylation close to the C-terminus. Immunofluorescence images using antibodies to ARO and RON4, suggested that the localization of rhoptries was impaired by IMP-1002 treatment. By homology with *T. gondii* ARO, the *N*-myristoylated ARO is involved in the correct positioning of rhoptries at the apical end of developing merozoites (Mueller, Samoo et al., 2016), and therefore NMT inhibition might lead to defective or mislocalized rhoptries. Such a mislocalization of rhoptries has been observed after parasite treatment with 2-bromopalmitate (2-BP) (Jones et al., 2012), which is a highly promiscuous inhibitor of lipid metabolism, with impacts on *S*-acylation (Davda, El Azzouny et al., 2013, Lanyon-Hogg, Faronato et al., 2017). ARO has two cysteines (Cys5 and Cys6) which are likely palmitoylated (Cabrera, Herrmann et al., 2012) and, together with the N-terminal myristoylation, involved in membrane anchoring and rhoptry positioning. These findings support the idea that loss of myristoylation of ARO changes the localization of the protein, and of the rhoptries away from the apical end of developing merozoites. RON4 is not *N*-myristoylated but is contained within rhoptries, and its mislocalization is consistent with the whole organelle being affected. In contrast to the effect on the rhoptries, IMP-1002 treatment during schizogony appeared to have no gross effect on either IMC formation or the localization of a micronemal marker.

The presence of IMP-1002 had a direct effect on protein *N*-myristoylation and abundance, as shown by two proteomic approaches that revealed differences in behaviour of some substrates following NMT inhibition compared with the DMSO control. Using chemical proteomics to examine proteins modified by YnMyr, *N*-myristoylation of sixteen proteins was significantly reduced during NMT inhibition compared with the DMSO-treated control. The YnMyr modified N-terminal peptide of several NMT substrates was also identified directly, including that of a metal-dependent protein phosphatase (PPM6) and a putative acylated pleckstrin-homology domain containing protein (APH), which had not been identified previously (Wright et al., 2014). Due to the co-translational nature of *N*-myristoylation, only those proteins synthesized during the YnMyr labelling window would have been purified and detected. Six of these substrates also showed a significant difference in protein abundance after IMP-1002 treatment compared with DMSO treatment, suggesting that NMT inhibition affects their overall stability. In addition to NMT substrates, other proteins were also decreased in abundance as a result of NMT inhibition. For example, IMP-1002 treatment led to a significant reduction of proteins involved in DNA chromatin organization and assembly, such as histones, and proteins involved in DNA replication, transcription and protein translation. A second large group included secreted and exported proteins, especially some targeted to the erythrocyte membrane. The export of parasite proteins to the erythrocyte is maximal in early erythrocyte asexual stages, but some proteins are also expressed in schizonts, stored in the apical organelles and then transferred to the erythrocyte at invasion (Marti, Baum et al., 2005). The observed changes in protein abundance may be due to a combination of altered transcription, protein synthesis and protein degradation rates.

During intraerythrocytic development, protein synthesis starts to accelerate at around 18 to 24 hours post invasion (Holder … Freeman, 1982), with the peak during schizogony. Total protein synthesis, as measured by metabolic incorporation of a methionine analogue, is not directly affected by NMTi (Wright et al., 2014), consistent with a selectivity and mode of action distinct from a direct effect on translation. However, since *N*-myristoylation is a co-translational process, it is possible that NMTi and protein synthesis inhibition may be connected through common essential downstream factors or pathways. In the whole proteome analysis, proteins involved in DNA synthesis, transcription, translation and chromatin organisation were less abundant following IMP-1002 NMT inhibition compared with DMSO treatment, strengthening the hypothesis of a connection between DNA and protein synthesis and NMT inhibition. NMT inhibition may also delay parasite development, even though there was no observed effect on nuclear division, slowing down essential processes during schizogony and leading to changes in protein abundance.

In addition to causing protein mislocalization as observed for ARO, inhibition of NMT may also result in the misfolding of its substrates, leading to their degradation and reduced abundance. For example, NMT inhibition leads to death through apoptosis of several cancerous cell types, potentially as a result of endoplasmic reticulum (ER) stress and an unfolded protein response (Thinon, Morales-Sanfrutos et al., 2016). Inhibition of NMT may also lead to an imbalance in favour of other N-terminal protein modifications (NPMs) such as *N*-α-acetylation (NAT) carried out by N-terminal acetyltransferase (NATs) (Starheim, Gevaert et al., 2012), or N-terminal ubiquitination (Ciechanover & Ben-Saadon, 2004, Timms et al., 2019). Usually these NPMs occur co-translationally through ribosome-associated protein biogenesis factors (RPBs) that interact with the ribosome and show a degree of competition in their binding (Giglione, Fieulaine et al., 2015). For example, there is some indication of competition between *N*-α-acetylation and *N*-myristoylation (Castrec et al., 2018, Utsumi, Sato et al., 2001). Additional experiments are necessary to investigate this further. In summary, treatment with NMTi results in mislocalization and reduced abundance of certain substrates that together may be responsible for the observed phenotype. An NMT inhibitor used to study myristoylation has an effect on many substrates simultaneously and the phenotype reflects the resultant pleiotropic consequences.

Because it is difficult to draw conclusions as to which of the myristoylation provides the greatest contribution to the observed phenotypes, we supplemented the inhibitor studies with genetic approaches. We expected that proteins with the greatest reduction in YnMyr labelling and abundance are those most affected by NMT inhibition, and these might contribute most to the observed phenotypes. In order to study *N*-myristoylation of particular substrates in isolation, the proteomic data sets were used to select six NMT substrates for a G2A substitution screen to determine the essentiality of the N-terminal glycine. To address individually the importance of myristoylation for these substrates, we developed a competition screen to study parasite viability following the integration of N-terminal glycine or N-terminal alanine constructs to repair a double strand break induced by CRIPSR-Cas9. This screen showed that for four out of six substrates the parasites preferentially incorporated the glycine codon at the second position with a ratio of greater than 80%, and for ARO, GAP45 and S9C nearly 100%. This strongly suggests the essentiality of myristoylation for at least four of the tested substrates. This approach does not allow phenotypic characterization of parasites lacking the N-terminal glycine in a specific substrate; therefore, GAP45 was selected for further analysis using a gene complementation approach to study the phenotype that results from the lack of myristoylation of this NMT substrate.

GAP45 is essential for glideosome assembly and erythrocyte invasion, as shown recently by using an inducible DiCre system to knockout the gene (Perrin et al., 2018). To complement this knockout, we placed the GAP45 gene in the Pfs47 gene locus, using two forms of the gene: one in which the second codon of the open reading frame encoded glycine, and a second in which the second codon encoded alanine. Interestingly, in both these constructs GAP45 appeared to be correctly targeted within the cell and allowed assembly of the glideosome as indicated by the correct location of MyoA and MTIP, which are not present in the absence of the complementing GAP45 gene copy. Although only the WT and not the G2A protein was myristoylated, both the G2A and WT GAP45 proteins were palmitoylated to a similar extent. Therefore, the single point mutation in the GAP45 gene, resulting in the presence or absence of the N-terminal myristoylated glycine, had a profound effect on parasite invasion, indicating that GAP45 myristoylation is essential for the function of the motor in invasion but not for motor assembly. It is possible that a low affinity interaction between GAP45 and the PM is essential, but to facilitate the dynamic changes that may be necessary for motor function (such as the passage along the membrane of the moving junction between parasite and RBC) the strength of the interaction needs to be modulated by differential palmitoylation/depalmitoylation of the cysteine close to the N-terminus or by GAP45 interaction with other proteins. The requirement for GAP45 myristoylation in invasion is clear but further work is needed to clarify the importance of further mechanisms. For example, the role of Cys5 palmitoylation should be addressed in future experiments to investigate the necessity of a second modification of the protein to complement myristoylation for dynamic membrane binding (Peitzsch & McLaughlin, 1993).

In conclusion, using small molecule inhibitors of NMT and genetic methods to replace the N-terminal glycine in NMT substrates, we have shown the importance of these substrates and their myristoylation at different stages in parasite development. As a consequence of these multiple effects, inhibitors targeting NMT provide outstanding antimalarial parasite activity.

## MATERIALS AND METHODS

### Parasite culture

*P. falciparum* 3D7 parasites were cultured *in vitro* in RPMI 1640 medium containing 0.5 % (w/v) Albumax II at 2-5 % hematocrit as described (Trager & Jensen, 1976). Parasites cultures were gassed with 90 % N2, 5 % CO2 and 5 % O2 and incubated at 37 °C. Parasites were synchronized using 70 % Percoll gradients to purify schizont stages, with a subsequent reinvasion followed by sorbitol treatment as described (Knuepfer, Napiorkowska et al., 2017).

### Determination of parasitemia by flow cytometry

Synchronized parasites were incubated with DMSO or IMP-1002 and samples were fixed in 4% paraformaldehyde (PFA), 0.2 % glutaraldehyde for one hour at 45 h PI. Then, samples were washed in phosphate buffered saline (PBS) and labelled with 1:500 of 10mg/mL Hoechst 33342 (New England Biolabs, Cat# 4082S) for 10 min with a subsequent wash in PBS. For flow cytometry analysis, a BD CL1 Fortessa D Analyzer or Aria Fusion Sorter and Analyzer with FACSDiva software v8.0.1 were used with the 450-50 filter, counting 50,000 RBCs per sample. Data were analysed using FlowJo LLC 2006-2015. Gating for RBCs was achieved by plots of forward scatter area against side scatter area (gate = P1). Doublet discrimination required gating on a plot of forward scatter height against forward scatter width (gate = P2) followed by a plot of side scatter height against side scatter width (gate = P3). A Hoechst-stained uninfected RBC sample was used as a negative control to gate on the infected population only on a forward scanner area against UVA fluorescence with 450-50 standard filter (gate = P4). Parasitemia was determined by the number of cells identified in gate P4 as a percentage of those in gate P3. The median fluorescence intensity (MFI) of each sample was used to determine the median number of nuclei per sample by normalizing it to the MFI of a control sample containing synchronized rings with a known MFI corresponding to one nucleus.

### Subcellular fractionation

Parasites were subjected to sequential fractionation to determine the solubility of proteins, using a method described previously (Ezougou, Ben-Rached et al., 2014). Schizont proteins were fractionated by sequential solubilisation using hypotonic and high salt buffers to release soluble cytosolic proteins, followed by a high pH sodium carbonate extraction to solubilise peripheral membrane proteins (carbonate-soluble) but not tightly associated membrane proteins such as integral membrane proteins (carbonate-insoluble). The distribution of specific proteins in the different fractions was revealed by western blotting.

### Western blot analysis

Proteins separated by SDS-PAGE were transferred to nitrocellulose membrane using the iBLOT Transfer system (ThermoFisher Scientific). Following blocking overnight at 4°C in 5 % (w/v) dried milk, 0.05% (v/v) Tween20 in PBS (PBS-T), membranes were incubated with primary antibody for 1 hr at RT in 5 % milk in PBS-T, followed by three 5 min washes in PBS-T and a subsequent incubation with species-specific secondary antibody (goat-anti-rabbit/rat/mouse IgG-HRP, Invitrogen 1:2500) for 1 h in 5 % milk in PBS-T. After a final three 5 min washes, the membrane was incubated with either 1 ml of Amersham ECL substrate western blotting detection reagent (GE Healthcare lot# 9622301) or for higher sensitivity, BioRad Clarity Western ECL substrate (Cat # 170-5060), used according to the manufacturers’ instructions. The signal was visualized on a BioRad ChemiDoc MP Imaging System.

### Indirect immunofluorescence assay (IFA)

For IFA, thin smears of parasitized RBC on slides were air dried, fixed in 4 % PFA in PBS for 10 to 20 min, permeablized in 0.1 % (v/v) Triton X-100 in PBS for 10 min, and blocked with 3 % bovine serum albumin (BSA) in PBS for at least 30 min at 4 °C. Slides were then probed with the appropriate dilution of primary antibody in a humidified chamber at room temperature (RT) for 1 h before being washed three times in PBS. Secondary antibody conjugated with Alexa Fluor 488 or 594 was added for one hour at the appropriate dilution, followed by three washes in PBS. Slides were mounted in ProLong® Gold Antifade mounting medium containing DAPI (4’,6-diamidino-2-phenylindole), and viewed on a Nikon Eclipse Ni-E imaging system with a Hamamatsu Orca-flash 4.0 digital camera and a Plan apo λ 100x/1.45 oil immersion objective. Images were captured using Nikon NIS-Elements software, generating Z-stack images of individual parasites, using deconvolution options and exporting the image as a tiff file. Alternatively, images were processed using Fiji software (Schindelin, Arganda-Carreras et al., 2012). Identical exposure conditions were used for each wavelength in treated (rapamycin or IMP-1002) and control (DMSO) samples.

### Metabolic tagging of parasites in the presence or absence of IMP-1002

For YnMyr (also known as YnC12 or Alk-14; tetradec-13-ynoic acid) tagging experiments, purified parasites were labelled metabolically using 25 μM YnMyr added to the culture medium. For YnPal (also known as YnC14 or Alk-16; heptadec-17-ynoic acid) labelling, the compound was stabilized by base treatment and absorbed to BSA to maximize its uptake and incorporation (Thinon, Fernandez et al., 2018). The required amount (for example, 120 μl of a 50 mM stock of YnPal for 240 ml RPMI 1640) was combined with 600 μl of 0.01M NaOH and warmed to 70°C for 3 to 4 min, then 1.5 ml of warm 5 % BSA solution was added and the mix maintained at 37 °C for 3 to 4 min. The solution was added to the RPMI 1640 culture medium, filtered through a 0.2 μm filter, and then the parasites were fed with the YnCPal-containing medium. In all experiments, the final DMSO percentage did not exceed 0.05 %.

### Preparation of *P. falciparum* proteins and copper(I)-catalyzed alkyne-azide cycloaddition (CuAAC) labelling

Parasites of the appropriate stage were either purified through Percoll, washed and pelleted or directly pelleted without purification. The cell pellet was lysed in 0.15 % saponin, using one and a half times the pellet volume for 10 min on ice. Following centrifugation, the pellet was washed further with PBS until the supernatant was free of hemoglobin and stored at −80°C until use. The pellet was thawed in ten times its volume of 1 % (v/v) Triton X-100, 0.1 % (w/v) SDS in PBS containing protease inhibitors but without EDTA, sonicated for 1 min and then left on ice for 20 min. Insoluble material was removed by centrifugation and the supernatant was snap frozen and stored at −80°C. The protein concentration of the lysate was measured using a Pierce^TM^ BCA Protein Assay Kit (23225, ThermoFisher Scientific) following the manufacturer’s instructions.

The lysate was adjusted to 1 mg/mL protein with PBS, and premixed click reagents [100 μM azido-TAMRA-biotin (AzTB) capture reagent, 1 mM CuSO4, 1 mM Tris(2-carboxyethyl)phosphine (TCEP), 100 μM Tris[(1-benzyl-1H-1,2,3-triazol-4-yl)methyl]amine (TBTA); mixed in the order stated and pre-incubated for 2 min] were added at the equivalent of 6 μl click reaction mix to 100 μl protein solution (Mousnier, Bell et al., 2018, Wright et al., 2014). The sample was vortexed for 1 h at RT and the reaction quenched by the addition of 10 mM EDTA. Protein was precipitated with 2 volumes of methanol, 0.5 volumes of chloroform and 1 volume of water. After centrifugation for 10 min at 17,000 *g*, the top methanol/water layer was removed and 0.5 ml ice-cold methanol was added prior to vortexing and sonication to break up and disperse the protein disc. The protein was collected by centrifugation (17,000 *g* for 10 min at 4 °C) and air-dried for ~15 min, then re-dissolved to 10 to 20 mg/ml in PBS containing 2 % SDS, 10 mM DTT, with vortexing for 15 – 30 min.

### Protein enrichment for immunoblot analysis

For analysis of affinity purified proteins by SDS PAGE, precipitated samples were enriched using 25 μl of Neutravidin Agarose Resin for lysate containing 150 μg of protein. The resin was pre-washed 3x with 0.2 % SDS in PBS followed by an enrichment of the labelled proteins from the lysate. Following the pull down for two hours at RT with shaking, the supernatant was removed, and the beads were washed three times with 0.2 % SDS in PBS. Proteins were eluted by treatment of the beads with SDS-PAGE sample loading buffer containing DTT (at a final 100 mM concentration) and boiling for 10 min. Following a centrifugation step to remove any insoluble material, supernatants were loaded on the gel.

### Affinity purification of labelled proteins and proteomic sample preparation

After click chemistry, precipitation, and dissolution in 2% SDS in PBS, samples were diluted with PBS to 1 mg/ml protein. For proteomic analysis, labelled proteins were first enriched. An agarose mixture comprised of one third Neutravidin Agarose resin and two thirds Pierce Control Agarose resin (ThermoFisher Scientific) was prepared to minimize contamination of samples with neutravidin from the beads, and 30 μl of this resin mixture was used to enrich labelled protein from lysate containing up to 300 μg protein. The resin mixture was pre-washed three times with 0.2% SDS in PBS using at least five times the bead volume, then the protein solution was incubated with the resin for two hours at RT, with shaking. The resin was washed sequentially three times with 5 to 10 volumes of 1% SDS in PBS, twice with 50 mM ammonium bicarbonate (AMBIC) containing 4 M urea, and a further three times with 50 mM AMBIC followed by sample processing as described previously (Broncel, Serwa et al., 2016). To improve detection of cysteine-containing peptides, thiols were reduced and alkylated; proteins were reduced with 10 mM DTT in 50 mM AMBIC for 30 min at 55 °C and alkylated with 10 mM iodoacetamide (IAA) in 50 mM AMBIC for 30 min at RT in the dark. Proteins were digested with trypsin overnight (0.12 μg Trypsin Gold [Promega UK Ltd, Cat. # V5280] for 300 μg protein). 1.5 % (v/v) trifluoroacetic acid (TFA; ThermoScientific Cat. #28902) was added to inactivate the trypsin and peptides were desalted using stop-and-go extraction (STAGE) tips and reverse phase C18 poly(styrenedivinylbenzene) polymer cation exchange (SDB-XC) membranes. The peptides were eluted in 79 % acetonitrile (MeCN)/21 % water and dried using a Speed Vac concentrator. Prior to liquid chromatography-tandem mass spectrometry (LC-MS/MS) analysis, samples were dissolved in 15 μl of 0.5 % TFA, 2 % MeCN in water using vortex, brief sonication and a final centrifugation step at 17,000 *g* for 10 min at 15 °C to remove insoluble material. Eleven μl of each sample was transferred to an autosampler-compatible vial.

### Global proteome analysis: Tandem Mass Tag (TMT) labelling of peptides and high pH reverse fractionation

Fifty microliters of lysate from parasites grown with or without IMP-1002 were treated with methanol:chloroform:water. Precipitated protein was washed with 200 μl methanol, collected by centrifugation (17,000 *g* for 10 min at 4°C) and solubilized in 20 μl 50 mM TEAB containing 0.2 % ProteaseMAXTM Surfactant (Promega UK Ltd, Cat. # V2071) for 1-2 h with vortex and occasional sonication. The samples were reduced with 5 mM DTT in 50 mM TEAB for 20 min at 56 °C and alkylated with 14.85 mM IAA in 50 mM TEAB for 30 min at RT in the dark. Samples were trypsin-treated (1.8 μg Trypsin Gold for 50 μg protein) in 0.05 % ProteaseMAX and then TFA was added to a final concentration of 0.5 %, and incubated for 5 min at RT, to inactivate the trypsin. Peptides were purified by STAGE-tip and reverse phase C18 SDB-XC membrane, with elution in 70 % MeCN and 30% water, and dried.

The TMT10plex Label Reagent Set (ThermoFisher Scientific, Cat # 90309) was used according to the manufacturer’s instructions. Immediately before use, the reagents were equilibrated to RT and dissolved in anhydrous MeCN. Peptides were dissolved in 25 μl 50 mM TEAB with sonication for 10 min, and then 0.2 mg TMT label reagent was added and each sample incubated for 1 h. To quench the reaction, 8 μl 5 % hydroxylamine were added to the sample and incubated for 15 min. A small quantity (about 5 %) of each sample was used to check the labelling through an initial liquid chromatography-mass spectrometry (LC-MS) analysis to determine the ratio of labelled reporter ions. Prior to mixing, the ratio was corrected for any differences in labelling efficiency. Samples were combined into one tube in equal amounts and peptides were initially separated by high pH reverse fractionation with a gradient step wise elution from 5 – 50 % MeCN to increase the proteome coverage, using the ThermoFisher Scientific kit (Cat # 84868) according to the manufacturer’s instructions. Each fraction was then dried and redissolved in 15 μl 0.1 % TFA to allow 10 μl per injection.

### Proteomic data acquisition and analysis

For the peptides from proteins labelled with YnMyr in the presence or absence of IMP-1002, data were acquired on a Q-Exactive Hybrid Quadrupole-Orbitrap mass spectrometer (Thermo Scientific) with a 120-minute acquisition time. Peptides were resolved chromatographically on an Ultimate 3000 RS-LC nano system (Thermo Scientific) using a 50 cm x 75 μm EASY-Spray^TM^ C18 column (Thermo Scientific) at a flow rate of 250 nl/min. The elution conditions comprised a gradient of solutions A (0.1 % aqueous formic acid [FA] in water) and B (0.1 % FA in MeCN) over 2 h. Via nano electrospray ionization, the eluent was introduced to the Q Exactive, which was operated in data-dependent mode using a survey scan of 350 – 1650 m/z at a resolution of 70,000. Up to 10 of the most abundant isotope patterns with 2+ charge or higher from the survey scan were selected with an isolation window of 2.0 m/z and fragmented by HCD with normalized collision energies of 25%. Subsequent scans were acquired at a resolution of 17,500 from m/z 200 - 2000.

For the whole proteome, analysis was performed on an Orbitrap Fusion Lumos Tribrid mass spectrometer (Thermo Scientific) with a 120-minute acquisition time. Peptides were resolved chromatographically on an Ultimate 3000 RS-LC-nano System (Thermo Scientific), using a 50 cm x 75 μm EASY-Spray C18 column (Thermo Scientific) at a flow rate of 300 nl/min. The elution conditions comprised a gradient starting at 2 % B (0.1 % FA, 80 % MeCN and water) and 98 % A (0.1 % FA in water) and increasing to 27.5 % B over 110 min followed by an increase to 40 % B over 10 min, and a final increase to 90 % B over 1 min. Via nano electrospray ionization, the eluent was introduced into the Orbitrap Fusion Lumos, which was operated in ‘TMT acquisition mode’ and peptides were analysed using a 375– 1500 m/z scan range using quadrupole isolation at 120,000 resolution for an ion at 200 m/z. Tandem mass spectra were first collected using the ion trap and fragmented using 35 % collisional induced dissociation (CID). A dynamic exclusion list was employed to prevent repeat sampling (repeat count of 2, repeat duration of 15 seconds, exclusion list size 100, and exclusion duration of 30 seconds).

### Proteome data analysis

Peptides identification and quantification were conducted using MaxQuant software (versions 1.5.3.8 for YnMyr labelling and label free quantitation, and 1.6.0.13 for the whole proteome with TMT quantitation) using the PlasmoDB-29_Plasmodium3D7_Annotated Protein database. All mass spectrometry ‘.raw’ files were loaded directly into the MaxQuant software. Protein intensity values were calculated based on the intensities of their corresponding peptides, and analyses of both LFQ (YnMyr labelling) and TMT (whole proteome) experiments in MaxQuant were performed using the built-in algorithms. Cysteine carbamidomethylation was selected as a fixed modification, and methionine oxidation and N-terminal acetylation as variable modifications. For the YnMyr labelling and purification experiment, myristoylation was set as a variable modification using a composition of C(22) H(37) N(7) O(4) with a monoisotopic mass of 463.2907 on any N-terminus. For enzyme digestion, trypsin was selected, which allows cleavage C-terminal of Arg and Lys residues and LysC which allows cleavage after Lys residues. Up to two missed cleavages were allowed. The false discovery rate (FDR) was set to 0.01 for peptides, proteins and sites. Other parameters were used as pre-set in the software. ’Unique and razor peptides’ mode was selected to allow identification and quantification of proteins in groups (razor peptides are uniquely assigned to protein groups and not to individual proteins), and all identifications were based on at least two unique peptides. The data were analysed using Perseus version 1.5.6.0, Microsoft Excel 2010 and GraphPad Prism version 8 for all experiments.

MS data were also processed with PEAKS X+, which as a default performs de novo peptide sequencing prior to database searches, in order to improve the accuracy of the results. The software also searches for common PTMs (PEAKS PTM) and point mutations (SPIDER). The data were searched against the same database used in MaxQuant analyses. Trypsin was selected for database searches. The maximal mass error was set to 5 ppm for precursor ions and 0.01 Da for product ions. Cysteine carbamidomethylation was set as fixed modification and methionine oxidation and myristoylation (463.2907 on any N-terminus) were set as variable modifications. The maximal number of modifications per peptide was set as three. The false discovery rate was set to 0.01 for peptides and a minimum of 1 unique peptide per protein was required.

### Generation of repair and Cas-9 plasmids for the G2A/G2G competition screen of ARO, CDPK1, GAP45, ISP3, S9C and TRP

For each locus a rescue plasmid was used with 200 base pair homology regions either side of the G2A mutation and guide sequence. The sequence between the G2A mutation and the guides was recodonized using the Codon Usage Table from PlasmoDB (Aurrecoechea, Brestelli et al., 2009). The tool on the ctegd.uga.edu/ website was used to determine two guides for each target gene based on close proximity to the G2A mutation, total score as calculated by use of an efficiency score (Doench, Hartenian et al., 2014) and the CRISPRRater (Labuhn, Adams et al., 2018). Each construct was flanked by unique restriction sites (SacI and SacII) not present in any of the constructs or the pMK-RQ kanamycin resistance plasmid from GeneArt, for linearization prior to transfection. For each construct a second plasmid contained the same homology arms and retained a codon for glycine at position 2 with a synonymous mutation from the endogenous sequence.

### Analysis of the G2A screen by DNA sequencing

The G2A screen was analysed by Illumina MiSeq. Parasite genomic DNA was extracted and specific integration-selective primers containing a MiSeq adapter sequence were used to amplify a 289-466 bp fragment (depending on construct) around the codon encoding G2A/G2G. The PCR was performed and samples were cleaned as recommended by the manufacturer for preparation of the 16S Metagenomic sequencing library (Part # 15044223). To increase the number of sequence reads per sample, the KAPA HyperPrep kit was used according to the manufacturer’s instructions to label the PCR fragments by ligating indices at each end creating a unique barcode for each sample. A nine bp sequence around the glycine or alanine codon was used to determine the ratio of integrated G2A versus G2G.

### Cloning of constructs and transfection of *P. falciparum*

The *gap45:ha3:loxP∷comp_gap45[G2A]* construct was generated through PCR, digest, ligation and cloning using the *gap45:ha3:loxP∷comp_gap45* construct and the same guide (Perrin et al., 2018), and cloned into the pDC2-cam-Cas9-U6-hDHFRyFCU-plasmid (Knuepfer et al., 2017, Lim, LaMonte et al., 2016, MacPherson … Scherf, 2015). Guide and rescue plasmids were paired and ethanol precipitated prior to transfection. *P. falciparum gap45:ha3:loxP* (B11 background) (Perrin et al., 2018) or 3D7 parasites were used. For transfection, mature schizonts were electroporated using the Amaxa 4D electroporator (Lonza) and the P3 Primary cell 4D Nucleofector X Kit L (Lonza) and program FP158 (Moon, Hall et al., 2013), with 60 μg of linearized rescue plasmid and 20 μg of the CRISPR/Cas9 plasmid carrying the respective guide RNA. Selections were carried out as recently described (Knuepfer et al., 2017); parasites were cultured in the presence of 2.5 nM WR99210 for five days to select for parasites with the Cas9/guide plasmid. Transfected parasites were detected after 22 days, and DNA integration was confirmed by PCR amplification. Parasites were then treated with 1 μM 5-fluorocytosine (Ancotil) to remove residual Cas9/guide plasmid and cloned by limiting dilution after 37 days (Rosario, 1981). Individual clones were then screened by PCR amplification to confirm integration of the required DNA sequence.

### Analysis of parasite growth and invasion

To analyse the growth of *gap45:ha3:loxP∷comp_gap45[G2A]* Clone 01 and Clone 02 as well as *gap45:ha3:loxP* and *gap45:ha3:loxP∷comp_gap45[WT]* parasites were adjusted to a parasitemia of 0.1 % and treated with rapamycin or DMSO. At the beginning of the assay (in cycle 0) and at 72 h (cycle one) and 120 h (cycle two) post invasion, when parasites were at a late ring/early trophozoite stage, samples were processed and analysed by flow cytometry. Each experiment was set up in triplicate, and these biological replicates were complemented with the use of the two clones, which served as an additional biological repeat.

The invasive capacity of genetically modified parasites (*gap45:ha3:loxP* / *gap45:ha3:loxP∷comp_gap45[WT]* and *gap45:ha3:loxP∷comp_gap45[G2A]* treated with either DMSO or rapamycin), and 3D7 parasites treated with either 140 nM IMP-1002 or DMSO, was measured using Percoll-purified synchronized mature schizonts added to RBC at 1% hematocrit and a parasitemia of 1 to 3%. Samples were fixed with 4% PFA and 0.02% glutaraldehyde at 0, 4, and 24 h later, enabling the percentage of newly formed ring-infected RBCs to be determined by Hoechst-staining and flow cytometry. Experiments were performed in triplicate with blood from three different donors. The data were analysed with FlowJo and GraphPad Prism software to determine the standard deviation and perform a t-test for statistical significance of differences between the samples.

### Acyl-PEG exchange (APE) analysis of protein thioesters

For Acyl-PEG exchange (APE) analysis, a parasite lysate (in 1% Triton X-100, 0.1% SDS, and EDTA-free protease inhibitor cocktail) was treated as described previously (Percher et al., 2016). Each lysate was adjusted to 2 mg/ml protein and 92.5 μl of samples per condition were treated to reduce Cys residues with 5 μl neutralized TCEP at a final concentration of 10 mM for 30 min with nutation. These free Cys residues were then blocked by alkylation with 2.5 μl of *N*-ethylmaleimide (NEM) (freshly prepared 1 M solution, diluted to 25 mM final concentration). The reaction was stopped by protein precipitation using methanol-chloroform-water (4:1.5:3) with sequential addition of 400 μl methanol, 150 μl chloroform and 300 μl water (all pre-chilled on ice). Following centrifugation (20,000 *g* for 5 min at 4 °C), the methanol/aqueous layer was removed, 1 ml pre-chilled methanol was added, and after mixing, the protein was pelleted by centrifugation at 20,000 *g* for 3 min at 4 °C. The pellet was washed again with pre-chilled methanol and dried under vacuum (Centrivap Concentrator, Labconco). To ensure complete removal of NEM from the protein pellet, each sample was resuspended in 100 μl 50 mM triethanolamine, pH 7.3, 150 mM NaCl containing 1× protease inhibitor mixture (Roche), 5 mM PMSF (Sigma), 5 mM EDTA (Fischer), and 1,500 units/mL benzonase (TEA buffer)(Percher et al., 2016), containing 4 % SDS, warmed to 37 °C for 10 min, and briefly (∼5 sec) sonicated (Ultrasonic Cleaner, VWR), with two additional rounds of methanol-chloroform-water precipitation.

For hydroxylamine (NH2OH) cleavage of palmitoyl thioester bonds and subsequent alkylation of the cysteines with methoxy(polyethylene glycol)-maleimide (mPEG-Mal), the protein pellet was redissolved in 100 μl TEA buffer containing 4 % SDS, 4 mM EDTA and split into two 50 μl samples. One sample was treated with 150 μl TEA buffer pH 7.3, containing 0.2 % Triton X-100 and 3 M NH2OH at a final concentration of 0.75 M NH2OH. The second control sample was not treated with NH2OH but diluted with 150 μl of TEA buffer, 0.2 % Triton X-100. After incubation at RT for 1 h with nutation, the protein was precipitated with methanol-chloroform-water and redissolved in 100 μl TEA containing 4 % SDS, 4 mM EDTA, warmed to 37 °C for 10 min, and briefly (∼5 s) sonicated. Next, to each sample was added 150 μl TEA buffer containing 0.2 % Triton X-100 and 4 mM mPEG-Mal (10 kDa; Sigma) for a final concentration of 1 mM mPEG-Mal. Samples were incubated for 2 h at RT with nutation before a final methanol-chloroform-water precipitation. The protein precipitate was re-dissolved as described above, and samples containing 10 μg protein were resolved by 3 to 12 % gradient Bis-Tris SDS-PAGE. and analysed by Western blot using rabbit anti-GAP45 and anti-CDPK1 antibodies.

## ACKNOWLEDGEMENTS

We thank Robert Goldstone, Matthew Winder and Laura Cubitt of the Advanced Sequencing Facility, The Francis Crick Institute, for their help. ACS was a Francis Crick Institute/Imperial College London PhD student, supported by the Department of Chemistry, Imperial College London and The Francis Crick Institute. JMS was supported by the European Union Framework Programme 7 (Marie Curie Intra European Fellowship). EWT was supported by the Cancer Research UK Programme Foundation Award C29637/A20183. This work was supported by funding from The Francis Crick Institute (https://www.crick.ac.uk; FC001097), which receives its core funding from Cancer Research UK (FC001097), the UK Medical Research Council (FC001097) and the Wellcome Trust (FC001097).

## Database deposition

The mass spectrometry proteomics data have been deposited to the ProteomeXchange Consortium via the PRIDE partner repository (Perez-Riverol, Csordas et al., 2019) with the dataset identifiers PXD022704 and PXD021879.

Reviewer account details: **Username**, **Password**:Gz9UqXto; **Username**, **Password:**1×7nHgnd

