## Supplementary Tables and Figures for "Inhibition of protein N-myristoylation blocks *Plasmodium falciparum* intraerythrocytic development, egress, and invasion"

Figure S1

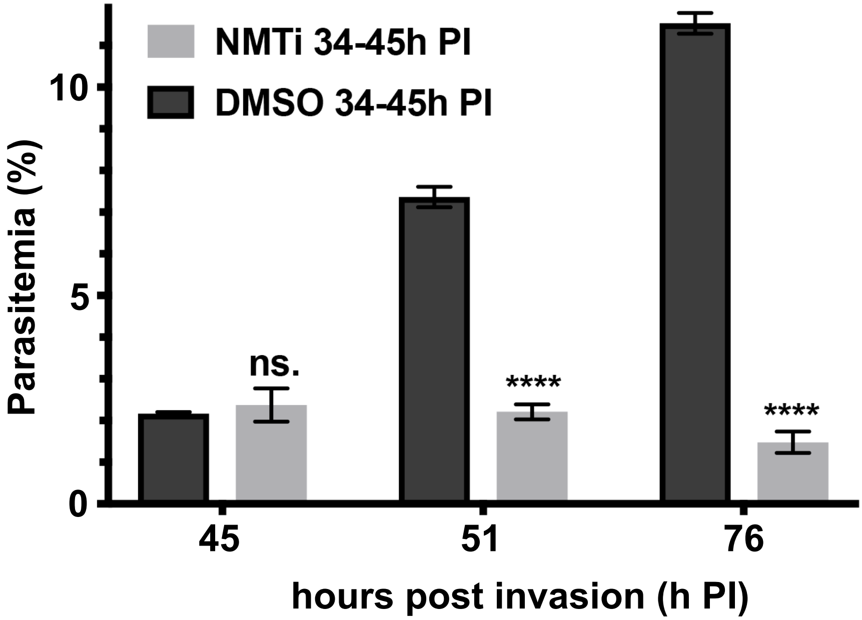

NMTi 34-45h PI
DMSO 34-45h PI
Parasitemia (%)
10
5
0
ns.
****
****
45
51
76
hours post invasion (h PI)

**Supplementary Figure 1. Inhibition of NMT during schizogony leads to a block in parasite development.** Parasitemia was significantly reduced in IMP-1002-treated culture compared with DMSO-treated control. Parasites were treated with IMP-1002 or DMSO from 34 to 45 hours post invasion (h PI). At the first sign of egress in the DMSO control (at 45 h PI), the growth medium was exchanged to drug-free medium and the parasites were quantified by flow cytometry six (51 h PI) and 31 (76 h PI) hours later. The difference in parasitemia was significant ($p < 0.0001$ for both 51 h PI and 76 h PI; unpaired Student t-test with Welch's correction not assuming an equal SD for each time point individually, $n = 3$).

Figure S2

**A. Modified N-terminal peptide of metal-dependent protein phosphatase PPM6 (PF3D7_1309200).**

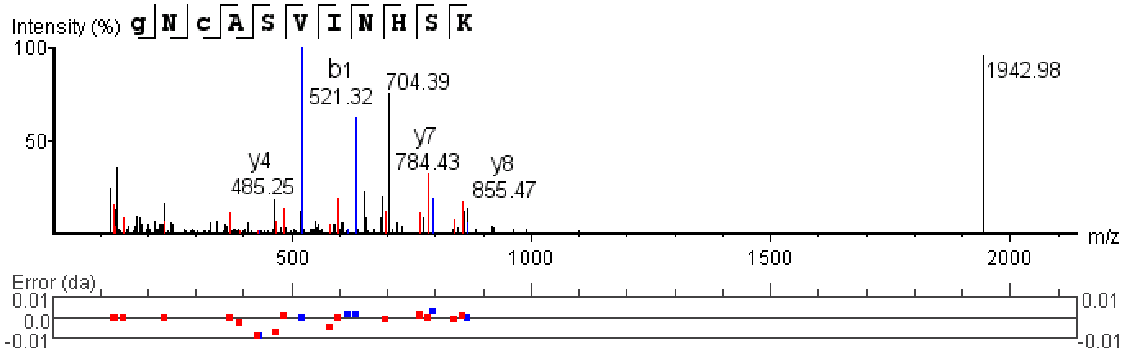


**B. Modified N-terminal peptide of acylated pleckstrin-homology domain containing protein (PF3D7_0414600).**

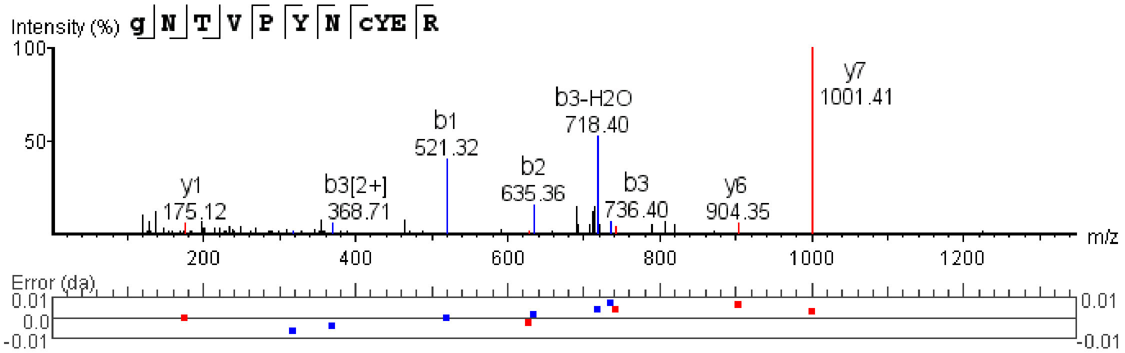

**Supplementary Figure 2. Direct identification of the modified N-terminal glycine of two NMT substrates by mass spectroscopy.**

Modified N-terminal peptides from A. Metal-dependent protein phosphatase 6 (PF3D7_1309200) and B. Putative acylated pleckstrin-homology domain containing protein (APH). The N-terminal modification and the peptide sequence is deduced from the parent ion mass and fragmentation pattern. The b1 ion (521.32, which correspond to the N-terminal glycine modified with YnMyr) is diagnostic of the metabolic incorporation of YnMyr by NMT into the protein.

Figure S3

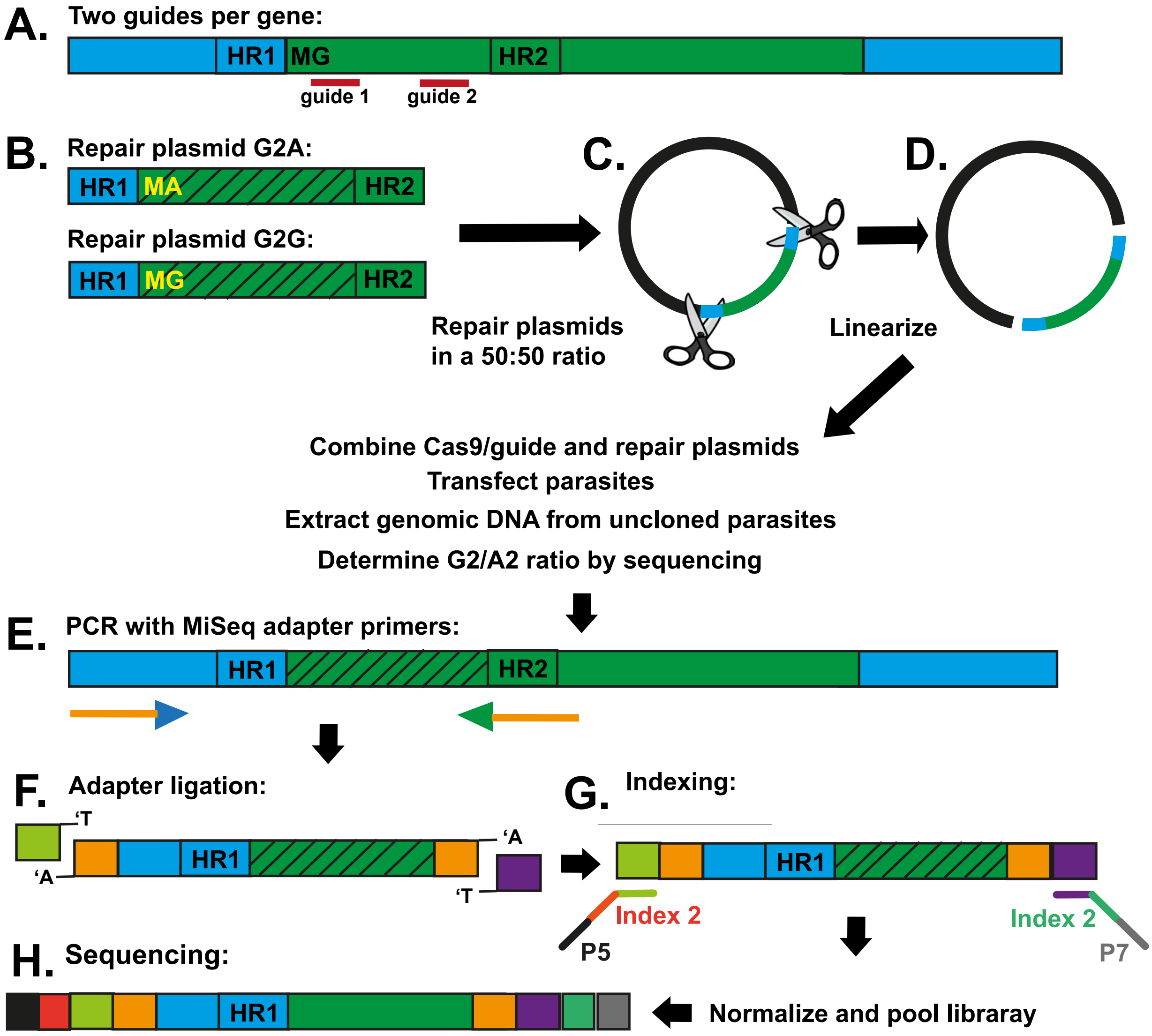


| Reads | GAP45 | ARO | CDPK1 | S9C | TRP | ISP3 |
|---|---|---|---|---|---|---|
| Expt. 1 | 2267 | 2012 | 43 | 1452 | 2327 | 660 |
| Expt. 2 | 2184 | 2003 | 8 | 1809 | 2003 | 573 |
| Expt. 3 | 2403 | | | | | |

**Supplementary Figure 3. A G2G/G2A CRISPR/Cas9-mediated viability screen**

**A.** For each gene of interest (ARO, CDPK1, GAP45, ISP3, S9C and TRP) two guide sequences were selected, except for *aro* for which the guide design was limited to one because the first exon was too short. **B.** Two repair plasmids with either a G2A point mutation or a silent G2G mutation were generated. **C.** The plasmids were mixed at a 50:50 ratio before linearization in the sequence flanking both homology arms (**D.**). Integration was facilitated by CRISPR-Cas9, and then following successful transfection, parasite genomic DNA was extracted. **E.** Integration specific primers allowed a selective PCR amplification of the integrated fragment, to which MiSeq adapter sequences were attached. **F.** The Illumina adapters were attached by ligation using the KAPA HyperPrep Kit. **G.** To discriminate between samples, the adapter-ligated fragments were labelled with indices by indexing PCR with two indices at each end creating a unique barcode. This step also added the attachment site for the MiSeq instrument (P5 or P7). **H.** Following Illumina sequencing, the ratio of the number of sequences for either the G2A or G2G variant provided an indication of the viability of each variant. The number of total reads with either the G2A or G2G sequence, for either two or three experiments, is indicated.

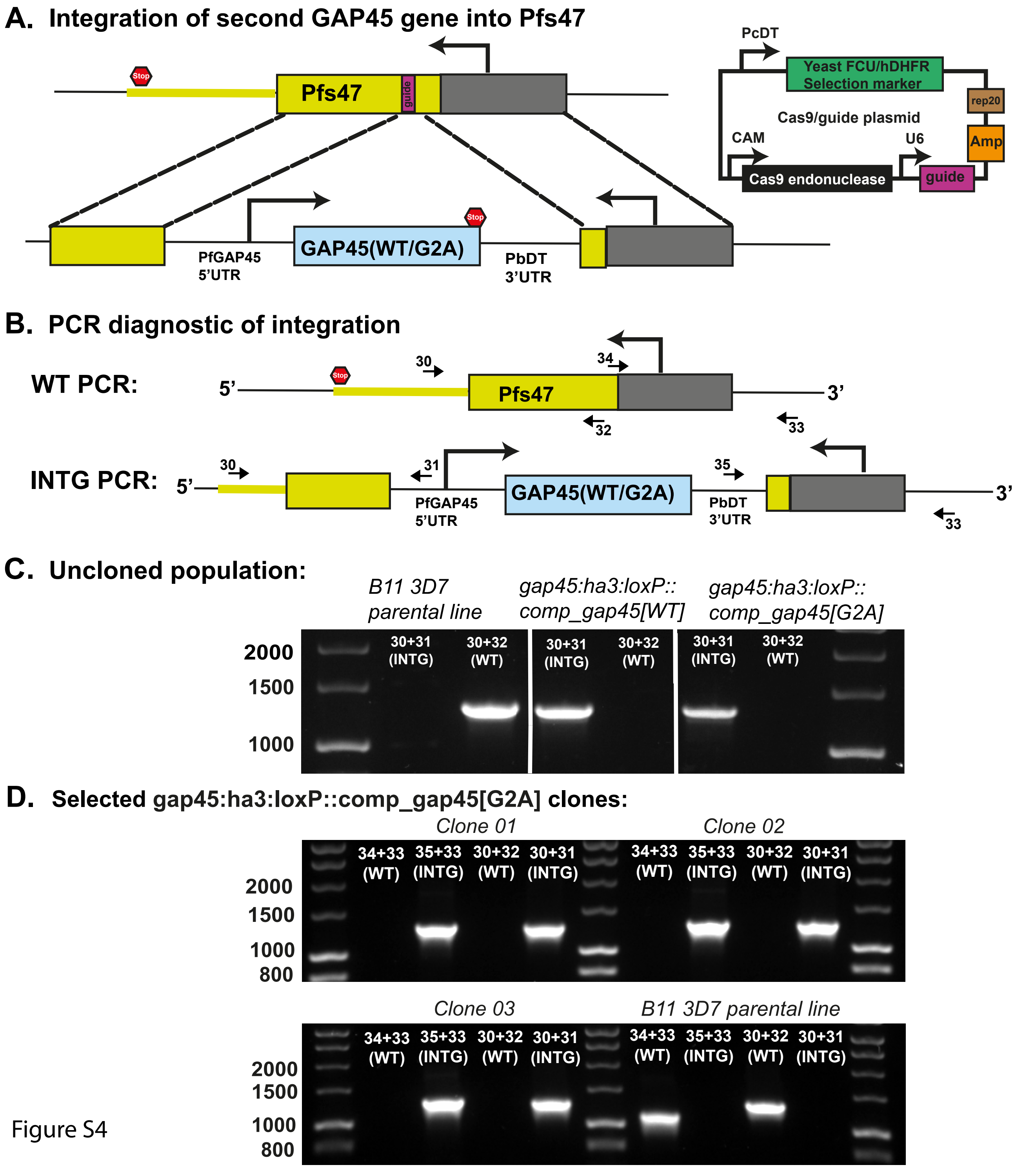
A. Integration of second GAP45 gene into Pfs47
Stop
Pfs47
guide
PcDT
Yeast FCU/hDHFR Selection marker
rep20
Cas9/guide plasmid
CAM
U6
Amp
Cas9 endonuclease
guide
GAP45(WT/G2A)
PfGAP45 5'UTR
PbDT 3'UTR
B. PCR diagnostic of integration
WT PCR:
5'
3'
30
34
32
33
INTG PCR:
31
35
C. Uncloned population:
B11 3D7 parental line
gap45:ha3:loxP::comp_gap45[WT]
gap45:ha3:loxP::comp_gap45[G2A]
2000
1500
1000
30+31 (INTG)
30+32 (WT)
D. Selected gap45:ha3:loxP::comp_gap45[G2A] clones:
Clone 01
Clone 02
Clone 03
B11 3D7 parental line
34+33 (WT)
35+33 (INTG)
800

Figure S4

**Supplementary Figure 4. Construction of a GAP45 gene complementation parasite line.**

**A.** The genetic complementation strategy used to introduce a second copy of the *gap45* gene coding for either an N-terminal glycine (G2G) or alanine (G2A) and its promoter sequence into the *pfs47* locus of a *gap45:ha3:loxP* parasite clone in the DiCre-expressing *P. falciparum* line B11, generating the *gap45:ha3:loxP::comp_gap45[WT]* and *gap45:ha3:loxP::comp_gap45[G2A]* lines. **B.** Schematic representation of the Pf47 locus with and without integration of the *gap45* gene, and the oligonucleotide primers within the 5' and 3' UTR regions used to analyse integration by PCR. **C.** PCR analysis of *gap45:ha3:loxP::comp_gap45[WT]* and *gap45:ha3:loxP::comp_gap45[G2A]* uncloned parasites showing 5' and 3' UTR integration. The B11 (3D7) line was used as a control for the WT parasite. **D.** PCR analysis of three selected *gap45:ha3:loxP::comp_gap45[G2A]* clones. The B11 3D7 line was used as a control for the WT parasite.

Figure S5

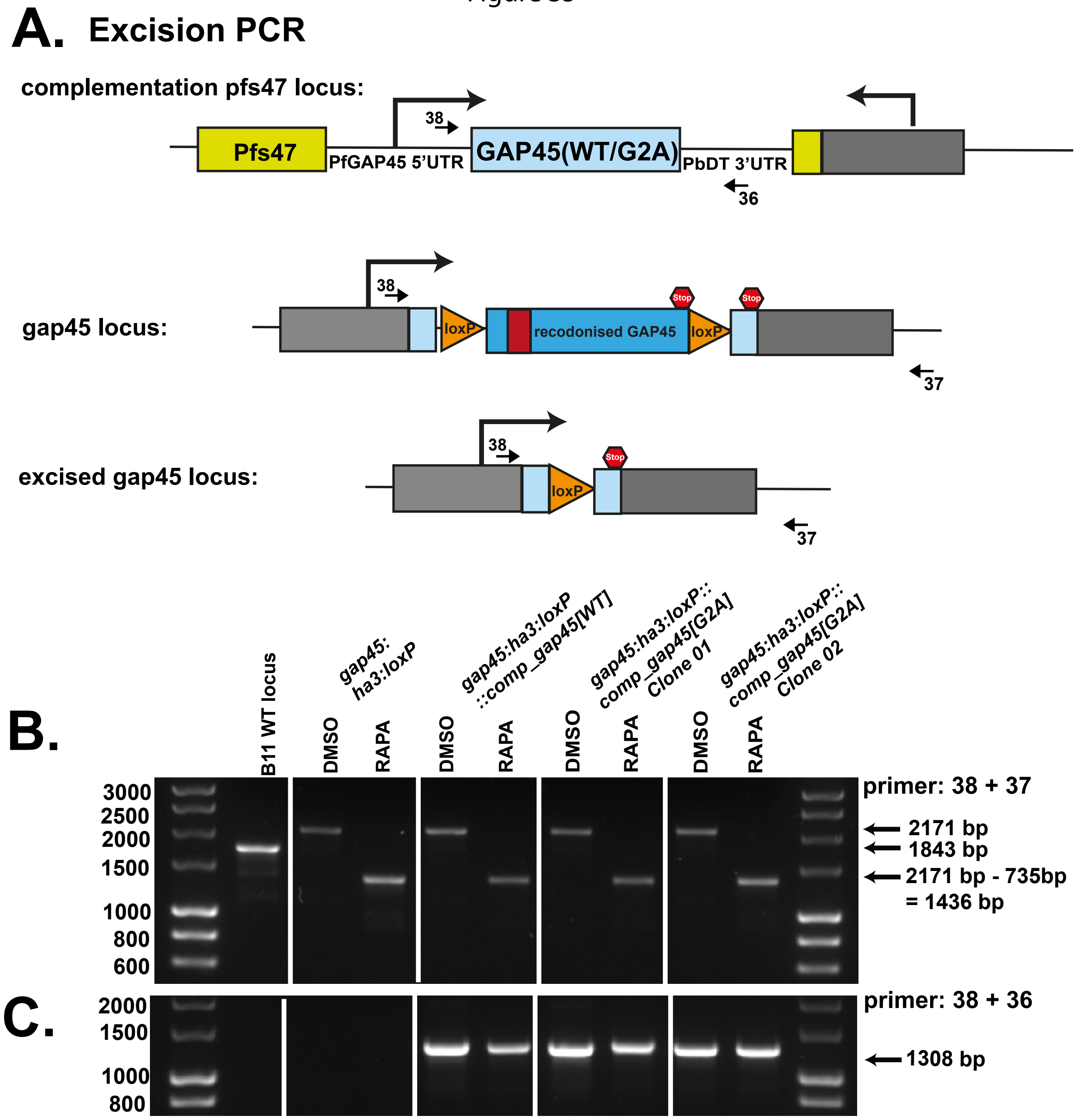
A. Excision PCR
complementation pfs47 locus:
Pfs47
38
PfGAP45 5'UTR
GAP45(WT/G2A)
PbDT 3'UTR
36
gap45 locus:
38
loxP
recodonised GAP45
loxP
Stop
Stop
37
excised gap45 locus:
38
loxP
Stop
37
B.
B11 WT locus
gap45:
ha3:loxP
gap45:ha3:loxP
::comp_gap45[WT]
gap45:ha3:loxP::
comp_gap45[G2A]
Clone 01
gap45:ha3:loxP::
comp_gap45[G2A]
Clone 02
DMSO
RAPA
DMSO
RAPA
DMSO
RAPA
DMSO
RAPA
3000
2500
2000
1500
1000
800
600
primer: 38 + 37
2171 bp
1843 bp
2171 bp - 735bp
= 1436 bp
C.
2000
1500
1000
800
primer: 38 + 36
1308 bp

**Supplementary Figure 5. Analysis of gene excision by PCR in *gap45:ha3:loxP, gap45:ha3:loxP::comp_gap45[WT]* and *gap45:ha3:loxP::comp_gap45[G2A]* parasite clones**.

**A.** Schematic representation of the complementation locus, the gap45 locus and the gap45 locus after rapamycin induced diCre-mediated excision, indicating the oligonucleotides used to analyse the parasites treated with rapamycin or DMSO by PCR, and oligonucleotides used to check presence of complemented construct after rapamycin treatment. **B.** Rapamycin induces excision at the *gap45* locus in all four parasite lines. **C.** Rapamycin has no effect on the complementation locus containing either the *gap45[WT] or gap45[G2A]* genes.

**Supplementary Table 1. Guide RNA sequences**

| Guide name | Sequence |
|---|---|
| ARO_G2A_Guide-01_F | ATTGATGGGAAATAATTGCTGTGC |
| ARO_G2A_Guide-01_R | AAACGCACAGCAATTATTTCCCAT |
| ARO_G2A_Guide-02_F | ATTGCTTCTGCTAAGTCTTGCTCA |
| ARO_G2A_Guide-02_R | AAACTGAGCAAGACTTAGCAGAAG |
| CDPK1_Guide_01_F | ATTGAGGAGAAGTAAATTTACGAA |
| CDPK1_Guide_01_R | AAACTTCGTAAATTTACTTCTCCT |
| CDPK1_Guide_02_F | ATTGTTACGAATGGAAATAATTAT |
| CDPK1_Guide_02_R | AAACATAATTATTTCCATTCGTAA |
| GAP45_Guide_02_F | ATTGATGTTCAAGAAGCAAAGTAA |
| GAP45_Guide_02_R | AAACTTACTTTGCTTCTTGAACAT |
| GAP45_Guide_03_R | AAACAACGTAAAGATATTGATGAA |
| GAP45_Guide_03_F | ATTGTTCATCAATATCTTTACGTT |
| ISP3_Guide_01_F | ATTGTTTCAGGACAATGAGATATA |
| ISP3_Guide_01_R | AAACTATATCTCATTGTCCTGAAA |
| ISP3_Guide_02_F | ATTGTAAGAGTGGAAATAGTATAA |
| ISP3_Guide_02_R | AAACTTATACTATTTCCACTCTTA |
| S9C_Guide_01_R | AAACACTAATCTTCAGACCGCACC |
| S9C_Guide_01_F | ATTGGGTGCGGTCTGAAGATTAGT |
| S9C_Guide_02_R | AAACCACCCACCTAGCTATTCAAA |
| S9C_Guide_02_F | ATTGTTTGAATAGCTAGGTGGGTG |
| TRP_Guide_01_F | ATTGTGCATTCGGAAGTAAGAATT |
| TRP_Guide_01_R | AAACAATTCTTACTTCCGAATGCA |
| TRP_Guide_02_F | ATTGTTATGCTAGCATGAAATTGT |
| TRP_Guide_02_R | AAACACAATTTCATGCTAGCATAA |

**Supplementary Table 2. Oligonucleotides used for PCR and sequence analysis**

| Name | Sequence |
|---|---|
| ARO_Intg_F1 | TATTATACCGTTGCCTTTCAAATGGCGGG |
| ARO_Intg_R1 | CATACATCTCTACCGGCGCAACAG |
| GAP45_Intg_F1 | aaacgtatggaagtgtaaaggg |
| GAP45_Intg_R1 | CTCGTCTATGTCCTTTCTCTTTGGC |
| ISP3_Intg_F1 | GGAAAAAGTTTTATGTTTCAGTCTGAATATTACC |
| ISP3_Intg_R1 | CTTTAATTGAGTTACCTGATTTGTACTTGTTAATCC |
| S9C_Intg_F1 | gaaatataaatacatgttaacacatagttataagataatacc |
| S9C_Intg_R1 | TTACTGTAACTTGGAGGATGTGGCC |
| TRP_Intg_F1 | GAAAATACATTTCTTCTTAATTCATATTG |
| TRP_Intg_R2 | CCACTTCTAATAACTTCATACTTGCG |
| CDPK1_Intg_F1 | cattttgatggtggcacttgcctttttgagg |
| CDPK1_Intg_R1 | ACCGTAGTTGTTACCGTTTGTGAACTTTGACC |
| CDPK1_WT_R1 | CCATTCGTAAATTTACTTCTCCTCG |
| ARO_WT_R1 | ATCTCTTCCTGCACAGCAATTATTTC |
| GAP45_WT_R1 | TTCATCAATATCTTTACGTTTGGGTTCC |
| ISP3_WT_R1 | CTGAAATATATCTATATTTGATTTACTG |
| S9C_WT_R1 | CTTTGAATAGCTAGGTGGGTGCGGTCTG |
| TRP_WT_R1 | TTTCATGCTAGCATAATTGTAATACTCC |
| ARO_MiSeq_F1 | TCGTCGGCAGCGTCAGATGTGTATAAGAGACAGTATTATACCGTTGCCTTTCAAATGGC |
| ARO_MiSeq_R1 | GTCTCGTGGGCTCGGAGATGTGTATAAGAGACAGCATACATCTCTACCGGCGCAACAG |
| GAP45_MiSeq_F1 | TCGTCGGCAGCGTCAGATGTGTATAAGAGACAGtaaacgtatggaagtgtaaaggg |
| GAP45_MiSeq_R1.2 | GTCTCGTGGGCTCGGAGATGTGTATAAGAGACAGCTCGTCTATGTCCTTTCTCTTTGGC |
| ISP3_MiSeq_F1 | TCGTCGGCAGCGTCAGATGTGTATAAGAGACAGGGAAAAAGTTTTATGTTTCAGTCTGAATATTACC |
| ISP3_MiSeq_R1 | GTCTCGTGGGCTCGGAGATGTGTATAAGAGACAGCTTTAATTGAGTTACCTGATTTGTACTTGTTAATCC |
| S9C_MiSeq_F1 | TCGTCGGCAGCGTCAGATGTGTATAAGAGACAGgaaatataaatacatgttaacacatagttataagataatacc |
| S9C_MiSeq_R1 | GTCTCGTGGGCTCGGAGATGTGTATAAGAGACAGTTACTGTAACTTGGAGGATGTGGCC |
| TRP_MiSeq_F1 | TCGTCGGCAGCGTCAGATGTGTATAAGAGACAGGAAAATACATTTCTTCTTAATTCATATTG |
| TRP_IMiSeq_R2 | GTCTCGTGGGCTCGGAGATGTGTATAAGAGACAGCCACTTCTAATAACTTCATACTTGCG |
| CDPK1_MiSeq_F1 | TCGTCGGCAGCGTCAGATGTGTATAAGAGACAGgaaaaataaaatcgtagaaaatgtttcccc |
| CDPK1_MiSeq_R1 | GTCTCGTGGGCTCGGAGATGTGTATAAGAGACAGGTAGTTGTTACCGTTTGTGAACTTTGACC |
| GAP45_MiSeq_F1.2 | TCGTCGGCAGCGTCAGATGTGTATAAGAGACAGtaaacgtatggaagtgtaaaggg |
| AJP_161 | gtgtatatttaccttacatttatctcc |
| AJP_162 | tatgacttgttcacttgctagtgtac |
| AJP_169 | gccaactgtagttgggtcatc |
| AJP_164 | cattcctaacacattatgtgtataaca |
| AS_GAP45C_Intg_WT3UTR | gtcaggggattaaaataaaaatc |
| AJP_163 | ATTGAGCAGAGGATATGCGCATAATGGT |
| G45_WTC_pbDT3_R | GCACACAACATACACATTTTTACAG |
| Integ_G45_3UTR_R | gatttcgatgaaattttaatttttttttaatc |
| AJP_93 | tgtttaatacatactgtgtaatcctt |

Upper  and lower case sequences are from exons and non-coding regions, respectively.

**Supplementary Table 3. Percentage of PEGylated and non-PEGylated protein in cell extracts measured by western blotting with anti-GAP45 and anti-CDPK1 antibodies (see Figure 7C).**

| **Protein** | | **Parasite line, % of protein signal** | | | |
|---|---|---|---|---|---|
| | | ***gap45:ha3:loxP::comp_gap45[WT]*** | | ***gap45:ha3:loxP::comp_gap45[G2A]*** | |
| | Rapamycin | - | + | - | + |
| **GAP45** | 2-PEG-protein | 13 | 8 | 13 | 15 |
| | 1-PEG-protein | 49 | 44 | 51 | 43 |
| | protein | 38 | 48 | 36 | 41 |
| **CDPK1** | 1-PEG-protein | 33 | 28 | 29 | 26 |
| | protein | 67 | 72 | 71 | 74 |

**Supplementary Data 1. All proteins from parasites metabolically labelled with YnMyr in presence or absence of IMP-1002 and bound and eluted from Neutravidin column**

**(Excel spreadsheet: Supplementary Data 1)**

**Supplementary Data 2. All proteins from parasites treated with IMP-1002 or DMSO**

**(Excel spreadsheet: Supplementary Data 2)**
